# A multivariate regressor of patterned dopamine release predicts relapse to cocaine

**DOI:** 10.1101/2022.10.31.514534

**Authors:** Miguel Á. Luján, Reana Young-Morrison, Sheila A. Engi, Brandon L. Oliver, Lan-Yuan Zhang, Jennifer M. Wenzel, Yulong Li, Natalie E. Zlebnik, Joseph F. Cheer

## Abstract

Drug addiction is characterized by a sustained vulnerability to relapse even after long periods of abstinence. A deeper understanding of the brain systems underlying this state could inform therapeutic strategies with novel prognostic biomarkers aimed at preventing renewed drug seeking. Most drugs of abuse, in particular psychostimulants such as cocaine, lead to long-lasting mesolimbic dopamine system adaptations, that ultimately facilitate drug seeking following exposure to drug-paired cues. This “dopaminergic hypothesis” of relapse has been previously addressed, but technical limitations in measuring *in vivo* dopamine release have precluded the assessment of its sufficiency without introducing pharmacological, electrical, or optogenetic confounds. Using a dopamine receptor-based fluorescent sensor in freely moving mice, we show that long-lasting dopamine recordings in the nucleus accumbens (NAc), throughout the animal’s entire history of cocaine self-administration, are strong predictors of relapse as well as the time it takes an animal to extinguish its drug seeking behavior. Moreover, we reveal previously unseen sex-specific trajectories of cocaine-related phasic dopamine responses from acquisition to relapse. We show that males exhibit higher-amplitude phasic dopamine responses, a trait accompanied by a greater resistance to extinguish their cocaine seeking, compared to females. Furthermore, we show that a semi-parametric model of the transition to extinction – using only multivariate patterns of dopamine release and sex as covariates – faithfully recapitulates male-specific vulnerability to persistent cocaine seeking. In conclusion, we present a predictive model of reinstatement behavior that uses information exclusively conveyed by NAc phasic dopamine responses, thus confirming, and actuating the sufficiency of the dopaminergic hypothesis of relapse.

## Introduction

Cocaine is the most consumed illicit psychostimulant in the world, with a resurgence of cocaine use disorders within the last decade^1^. It is estimated that 16% of the 18 million people that use cocaine will eventually become affected by a cocaine use disorder^2^. The increased risk of relapse associated with prolonged cocaine use is a major contributor to these sobering statistics. Given the limited efficacy of existing treatments, it is of capital importance to identify prospective biomarkers that can serve as accurate predictors of relapse to improve the diagnosis, prognosis, and treatment of cocaine use disorder^3^.

Clinical and preclinical evidence have outlined the neurobiological foundations of cocaine relapse^4,5^. Increased vulnerability to relapse results from maladaptive potentiation of the mesolimbic dopamine system, which remains compromised even after long abstinence periods^6,7^. The ventral tegmental area (VTA) is the originating point of what has been described as the “final common pathway” of reinstatement^8^. From there, dopaminergic projections to the nucleus accumbens (NAc), the medial prefrontal cortex, and the basolateral amygdala facilitate the initiation of drug-directed behaviors, ultimately leading to relapse^7–12^. Nevertheless, the brain circuitry associated with an increased risk of relapse is not solely related to enhanced dopamine dynamics^13^ as other non-dopaminergic motivation systems^14^ may determine relapse behavior independently of a common mesolimbic dopamine substrate^15^. However, recent experiments have re-emphasized the role of VTA dopamine neuron activation to promote relapse to reward-associated cues^16^. Technical limitations have so far precluded the formulation of an unambiguous “dopaminergic hypothesis” of relapse that does not introduce artificial manipulations to alter the mesolimbic dopamine system (e.g., optogenetics, chemogenetics, lesions, electrical stimulation, pharmacological agents, etc.). The recent emergence of dopamine-specific genetically-encoded fluorescent sensors bridges this gap in knowledge, by testing whether dopaminergic variables carry sufficient information to reproduce and predict cocaine relapse without artificially manipulating the anatomical substrate. The GrabDA and dLight families of sensors enable longitudinal recordings of subsecond dopamine dynamics in freely moving animals during extended periods of time^17,18^, a combination of features unmatched by voltammetric (compromised long-term stability) and microdialysis probes (low temporal and spatial resolution).

Similar temporal limitations have also hindered the ability to probe dopamine dynamics throughout the entire animal’s history of cocaine taking – from initial consumption to relapse. A fiber photometry study recently suggested that dopamine signatures of alcohol seeking are characteristic of the drug-intake stage^19^. Despite this promising evidence, it remains unclear at which stage sensitized dopaminergic signaling faithfully determines future enhanced risk to relapse. Similarly, potential sex differences in these longitudinal cocaine-evoked dopamine dynamics have yet to be elucidated^20^. This is of notable importance, since human and animal studies reveal sex differences at every phase of drug addiction^21,22^ (but also see Nicolas et al.^23^). Women escalate to compulsive drug-taking more rapidly than men, and report larger difficulties to maintain abstinence^24^. Men show higher rates of transition between cocaine abstinence and use states, despite similar levels of baseline consumption^25^. Structural and functional differences in the dopamine system – among other brain systems – of males and females may underlie these differences^26–28^. Surprisingly, no study has systematically explored sex-specific patterns of phasic dopamine release in cocaine self-administering rodents.

Here, we utilize GrabDA-based fiber photometry to uncover cocaine-evoked phasic dopamine responses in the NAc throughout the animal’s entire history of contact with the drug, from acquisition to reinstatement. To refine the “dopaminergic hypothesis” of cocaine relapse, we test whether cued reinstatement of cocaine seeking is reproduced from the information obtained from multivariate patterns of accumbal dopamine activity. Another correlate of enhanced risk of relapse after cocaine consumption – resistance to extinguish cocaine-seeking behavior – is also tested with our mathematical model. Furthermore, we uncover previously unseen sex-specific trajectories of cocaine-evoked dopamine responses underlying male vulnerability to persistent cocaine seeking.

## Results

### Protracted behavioral and dopaminergic responses to voluntary, intravenous cocaine infusions in mice

An optical fiber was implanted into the NAc core to probe for changes in fluorescence induced by GrabDA_2m_ (Figure 1A, Methods). Following fiber implantation but prior to commencing behavioral experiments, animals (N = 12) were catheterized in the right jugular vein to allow for voluntary intake of intravenous cocaine (0.5 mg/kg/inf). During 6-8 weeks of daily recordings, accumbal dopamine responses accompanying cocaine intake and drug-predictive cues were sampled throughout the acquisition, maintenance, extinction, and reinstatement of cocaine-seeking behavior. During acquisition, mice had to nosepoke an active porthole placed in one side of the chamber in order to receive a 2-s long cocaine infusion paired with a discrete compound cue (light + tone) (Figure 1B). Nosepokes on the inactive hole, located on the opposite side of the chamber wall, had no programmed consequences but served as an indication of generalization of motor behavior. Animals had no previous training experience before cocaine self-administration. Figure 1C displays averaged responses on the active and inactive portholes during the acquisition (fixed ratio 1 reinforcement schedule, or FR1) and maintenance (FR3) phases of the experiment. Group-averaged, peri-event time histograms (PETHs), comprising trials from each experimental phase, served to characterize each subject’s dopaminergic responses following cue onset (0-2 s) and cocaine delivery (2-30 s). Significant GrabDA_2m_ transients were determined using a bootstrapping confidence interval (CI) procedure (95% CI, 1000 boot-straps)^29^ (Methods). A different set of animals followed the same training protocol but were reinforced with a sucrose pellet. Their dopaminergic responses served as the ground truth to illustrate the identifying features of cocaine-evoked dopamine transients in the NAc. During the first FR1 sessions, cocaine infusions elicited increases in extracellular dopamine release that were remarkably different from those following a natural reinforcer (sucrose pellet) (Figure 1D). Dopamine release upon cue presentation and reward delivery was initially higher following a natural reward. Cocaine infusions instead elicited steady, prolonged increases lasting a minimum of 30-s, reflecting the accumulation of extracellular dopamine due to the pharmacological action of cocaine at NAc dopamine transporters (DAT)^30–32^. Figure S1 depicts the full extent of the GrabDA_2m_ transients throughout an entire self-administration session, along with the modeled quantities of cocaine brain concentrations^33^ (Methods). A broader examination reveals that cocaine-evoked phasic dopamine transients returned to baseline levels in the order of minutes. Surges in modeled cocaine brain concentrations followed the same temporal scale, as expected, but kept accumulating throughout the 2-h long sessions (Figure S1). The difference between rewards were maintained later into the maintenance phase, under a FR3 reinforcement schedule (Figure 1E). Mice that acquired self-administration behavior advanced to a progressive ratio (PR) test, an exponentially growing schedule of reinforcement (Figure 1F) used to parse the reinforcing efficiency of cocaine and other rewards in rodents^34,35^. Breaking points, number of infusions obtained, total nosepokes and cocaine-evoked dopamine transients (Figure 1 G-J) were registered for further analysis. Twenty-four h after PR, animals underwent subsequent extinction sessions (Figure 1K) until seeking on the cocaine-paired porthole ceased (extinction criteria: reduction in active nosepokes to >40% of FR3 average, during two consecutive extinction sessions). GrabDA_2m_ averaged traces from the first (early extinction) and last extinction session (late extinction) were obtained and plotted in Figures 1L and M, respectively. Bootstrap 95% CI procedure analyses of early and late extinction trials revealed a decrease in dopaminergic encoding of cocaine-seeking in the active porthole. After the last extinction session, mice underwent a cue-induced reinstatement test (Methods), where cue presentation (but not cocaine infusion) followed every nosepoke on the active hole. As shown in Figure 1N, reappearance of cocaine-associated light + tone cue elicited again a robust dopamine transient in the NAc. GrabDA_2m_ transients of inactive nosepokes are shown in Figure S2.

**Figure 1.**
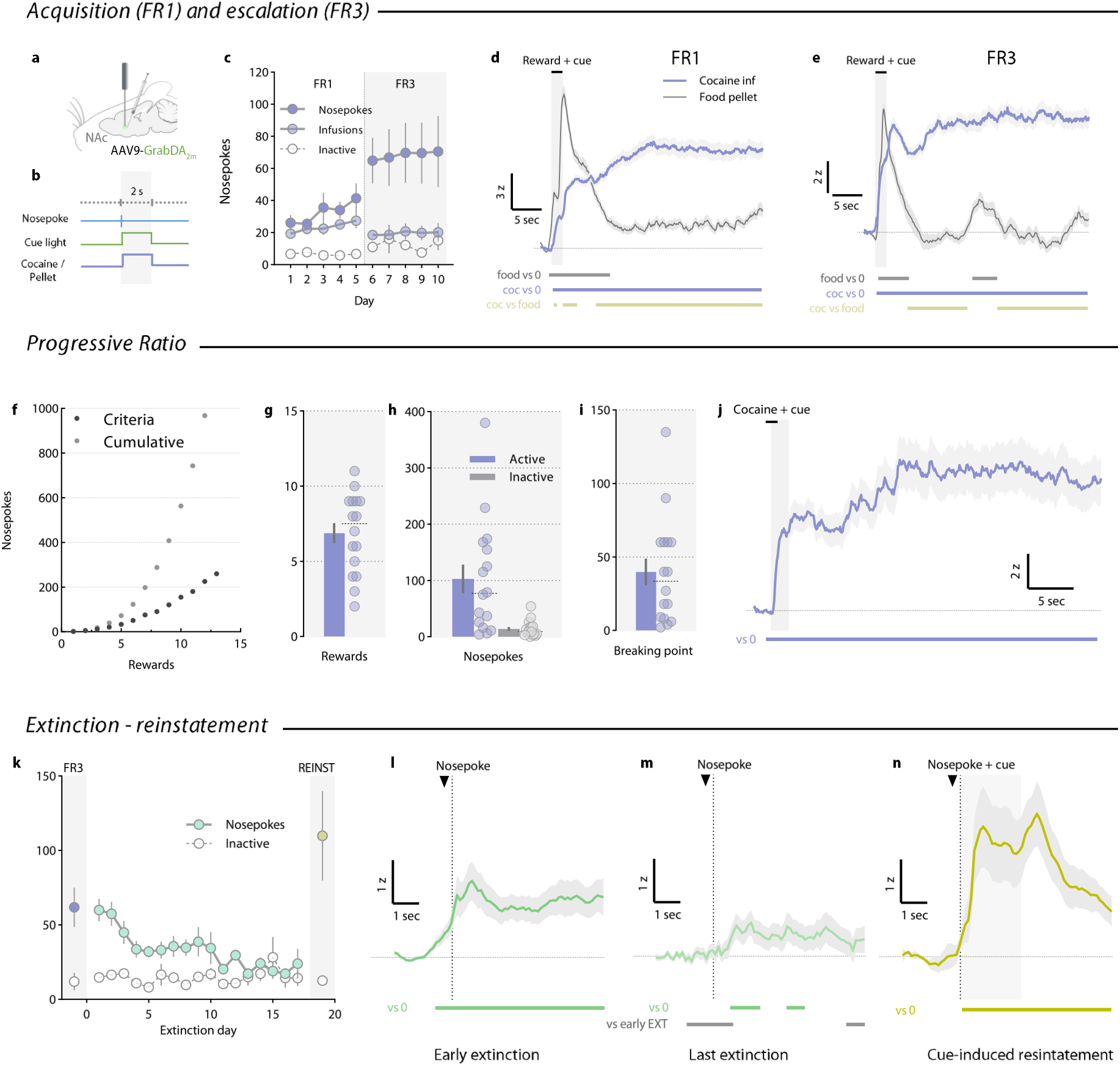
Phasic dopamine responses in the NAc during cocaine-seeking behavior in mice. **a**) GrabDA_2m_ was transduced in the NAc core, where an optical fiber was implanted. **b**) Mice were trained to nosepoke into a porthole in order to obtain a 2-s long cocaine infusion signaled by a contingent cue light and tone. **c**) Active and inactive nosepokes and rewards from the acquisition (FR1) and maintenance (FR3) phases of the cocaine/food self-administration (mean ± SEM). **d**) NAc GrabDA_2m_ transients (mean ± SEM) centered −2 s to +30 s around cue onset and obtained during FR1 sessions. Colored bars below traces represent periods significantly different from 0 or other traces, as defined by bootstrapped 95% confidence intervals. **e**) NAc GrabDA_2m_ transients (mean ± SEM) obtained during FR3 sessions. **f**) Schematic representation of the response requirement during the progressive ratio (PR) test. **g**) Number of rewards (mean ± SEM) obtained on the PR test. **h**) Total active/inactive nosepokes (mean ± SEM) from the PR session. **i**) Breaking points (last response requirement reached, mean ± SEM) of cocaine demand on PR. **j**) NAc GrabDA_2m_ transients (mean ± SEM) obtained during PR. **k**) Active/inactive nosepokes (mean ± SEM) during extinction and cue-induced reinstatement of cocaine-seeking behavior. FR3-averaged responding is shown for reference. 1-way repeated measures (RM) ANOVA reported a significant decrease of cocaine-seeking behavior across days (F_16,149_ = 4.05; *p* < 0.0001). l, m) NAc GrabDA_2m_ transients (mean ± SEM) from the first (**l**) and last (**m**) extinction sessions, centered −2 s to +5 s around every active nosepoke. **n**) NAc GrabDA_2m_ transients (mean ± SEM) from the cue-induced reinstatement session, centered −2 s to +5 s around cue onset.

### Predicting reinstatement of cocaine seeking from multivariate patterns of dopamine responses during each animal’s history of drug self-administration

Prior to the emergence of genetically encoded, fluorescent dopamine reporters^17,18^, *in vivo* dopamine measures were constrained either by long-term stability (i.e., voltammetry) or by temporal and spatial resolution (i.e., microdialysis). These technical limitations may have hindered our ability to understand the relevance of phasic, subsecond dopamine signatures of cocaine seeking in the context of long-term drug addiction. As a result, potential relationships between adaptations in drug-evoked dopamine responses and relapse to cocaine remain unexplored. Our first approach to uncover associations between cocaine-evoked phasic dopamine release and the reinstatement of cocaine-seeking behavior consisted of Pearson’s correlation matrices. Behavioral measurements (number of nosepokes or infusions during each phase, and number of days to extinction) were also correlated with the number of nosepokes on the reinstatement test. Figures 2A and 2B show the Pearson’s matrices resulting after correlating every dopaminergic (Figure 2A) or behavioral (Figure 2B) measurement obtained during the entire history of cocaine self-administration. Correlations with the number of active nosepokes during reinstatement are highlighted on both panels. As expected, there was a higher degree of correlation amongst behavioral measurements (Figure 2B). This reflects the notion that high-responding animals steadily maintain responding during the entirety of the self-administration experiment. The number of nosepokes emitted during FR1 and FR3, as well as the number of infusions, was significantly correlated with nosepokes during cued reinstatement. On the contrary, correlations between dopaminergic variables and reinstatement nosepokes were fewer and more diverse (Figure 2A). Interestingly, dopamine amplitudes (averaged GrabDA_2m_ Δ F/F_0_ z-scores) aligned to cue onset (0-2 s) during FR1 (r = 0.61, *p* = 0.047) and PR (r = 0.81, *p* = 0.003) (depicted in Figure S3) were positively correlated with the incidence of reinstatement. This result highlights the importance that early phasic dopamine responses may have in predicting later relapse to cocaine use. Following this line of evidence, we next implemented a multiple linear regression (MLR) model that would allow us to predict cue-induced reinstatement of cocaine-seeking behavior solely by using dopamine responses (GrabDA_2m_ transient amplitudes) observed throughout the entire history of cocaine self-administration for each animal.

**Figure 2.**
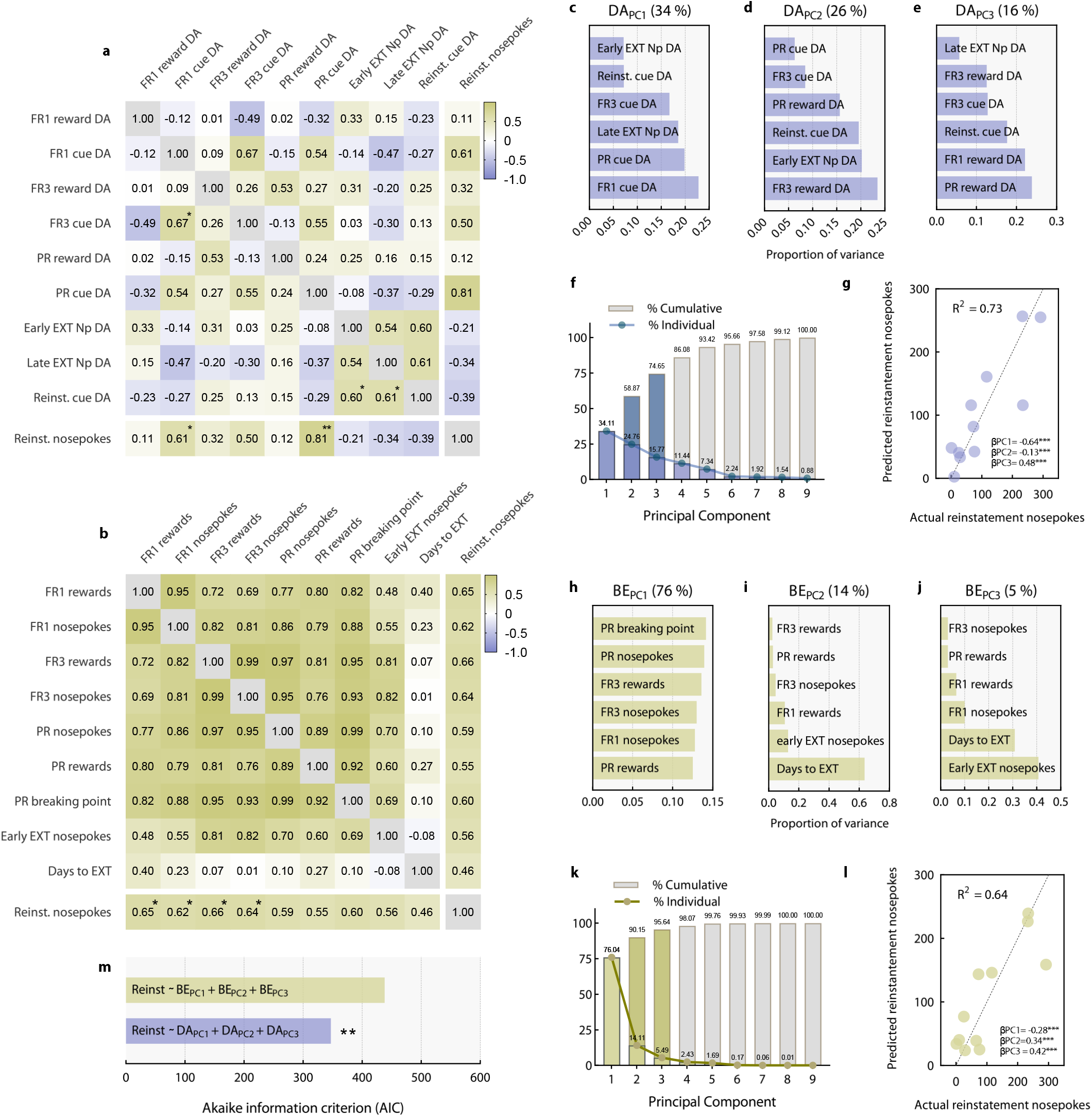
Low-dimensional dopamine signatures robustly predict reinstatement of cocaine-seeking behavior. **a**) Pearson’s correlation matrix obtained from GrabDA2m Δ F/F_0_ z-scores measurements made during cocaine self-administration. Person’s *r* values from each correlation are color-coded and shown within each cell. Nosepoke correlations observed during the relapse test are shown separately. Statistically significant correlations (\**p* < 0.05, \*\**p* < 0.01) are marked with asterisks. **b**) Pearson’s correlation matrix obtained from all behavioral measurements made throughout each cocaine self-administration session for all animals. For the sake of clarity, only significant nosepoke correlations (\**p* < 0.05) during reinstatement are shown. **c**-**e**) Proportion of variance explained by operationalized dopaminergic variables of each principal component (DA_PC_) obtained from principal component analysis (PCA). **f**) Percentage of total variance (%) explained by each of the DA_PC_ factors derived from PCA. Principal components selected for a multiple linear regression model (MLR) are colored in dark blue. **g**) Actual vs. predicted reinstatement nosepokes obtained from MLR with DA_PC1_, DA_PC2_, and DA_PC3_ as covariates. The model’s R^2^ and β-parameter estimates for each DA_PC_ are shown as an inset. Asterisks indicate β significantly different from 0 (\*\*\**p* < 0.0001). **h**-**j**) Proportion of variance explained by behavioral variables of each principal component (BE_PC_) obtained from PCA. **k**) Percentage of total variance (%) explained by each of the BE_PC_ factors derived from PCA. Principal components selected for the multiple linear regression model (MLR) are colored in yellow. **l**) Actual vs. predicted reinstatement nosepokes obtained from MLR with BE_PC1_,BE_PC2_,BE_PC3_ as covariates. The model’s R^2^ and β-parameter estimates for each BE are shown as an inset. Asterisks indicate β significantly different from 0 (\*\*\**p* < 0.0001). **m**) Model comparison using the Akaike information criteria (lower is more parsimonious) supports the superiority of the DA_PC_ model over the BE_PC_ model (ΔAIC = −90.24; \*\**p* = 0.006). “Reinst” = reinstatement, “Np” = nosepoke, “EXT” = extinction, “DA” = dopamine.

Given the number of independent variables available, we conducted a principal component regression (PCR) to circumvent the problem of overfitting our linear model with too many free parameters. This PCR method also allowed us to avoid biasing the model by arbitrarily selecting some dopamine variables and to reduce multicollinearity. In addition, PCR gave us access to low-dimensional features of the dopamine response space that may reflect hidden degrees of freedom. The resulting components obtained for the dopaminergic variables are shown in Figures 2C-E. The three first principal components, explaining near 75% of the total variance (Figure 2F), were selected for further analysis. The first principal component (DA_PC1_) explained 34% of the total variance and was exclusively composed of cue-evoked dopamine release variables, therefore reflecting a stable signature of accumbal dopamine responses to drug-paired cue presentation. Principal components 2 and 3 (DA_PC2_, DA_PC3_) yielded a more multifaceted combination of dopamine responses. The inclusion of cocaine-evoked dopamine variables (post-drug delivery, 2-30 s) among the top contributing parameters suggests a stable signature of reward-evoked dopamine responses. PCA factor loading and subject’s scores are shown in Figure S4. Next, DA_PC1_, DA_PC2_ and DA_PC3_ were used as estimators in a MLR model (hereafter referred to as DA_PC_) to predict the number of nosepokes observed during reinstatement (Methods). Our results indicate that multivariate patterns of cocaine-related phasic dopamine responses (DA_PC1_, DA_PC2_ and DA_PC3_) can robustly predict reinstatement behavior (R^2^ = 0.73) (Figure 2G). The model also revealed that all DA_PCs_ contributed to the observed variance during reinstatement, suggesting that cue- and cocaine-aligned dopamine transients are critical gatekeepers of cued reinstatement to cocaine-seeking behavior. Considering the relatively small sample used, we sought to replicate these findings with an alternative approach. Using Bayesian Poisson regression (Methods), we confirmed the model’s prediction by showing that the posterior estimates of DA_PC1_ (β = −0.65, lCI = −0.71, uCI = −0.59; 95%), DA_PC2_ (β = −0.13, lCI = −0.18, uCI = −0.08; 95%) and DA_PC3_ (β = 0.48, lCI = 0.44, uCI = 0.55; 95%) β coefficients significantly differed from 0 and matched the previous MLR model. Figure S5 shows the high concordance between the observed reinstatement data and the maximum *posterior* predicted distributions of the Bayesian model.

To examine how informative these low-dimensional dopamine signatures can be, we then compared the DA_PC_ model against another PCR model obtained from the behavioral measurements, as similarly seen in Flagel et al.^36^. Given the high degree of covariance between the behavioral variables, such a behavioral PCR should yield a reasonably good predictive model to compare. It is also safe to assume that nose-poking behavioral variables would be the best predictors of another nose-poking behavioral variable. To keep the number of free parameters consistent among models, we selected the three first components resulting from the initial PCA (Figure 2H-J), which explained near 97% of the behavioral variance (Figure 2I). The first behavioral principal component (BE_PC1_; 76% of the explained variance) was exclusively composed of number of nosepokes and infusions during sessions in which cocaine was available, thus reflecting cocaine reinforcement. The second principal component (BE_PC2_) was predominantly defined by the time that animals took to reach extinction criteria. The amount of nosepokes given when access to cocaine was initially removed (craving) featured the third principal component (BE_PC3_). PCR yielded a considerably good prediction of reinstatement behavior (R^2^ = 0.64), to which all principal components contributed. Using the Akaike information criteria (AIC), we found that the DA_PC_ model fit reinstatement behavior data better than the BE_PC_ model (ΔAIC = −90.24; *p* = 0.006) (Figure 2M). Furthermore, we performed additional PCR analyses to illustrate that the superiority of the DA_PC_ model did not depend on the method used to select principal components (Figure S6). Finally, we iteratively tested how well the DA_PC_ model predicted other behavioral outcomes (such as FR1/3 responding, PR breaking points, etc.). A comparison between the percentage of variance explained (R^2^) of each dependent variable indicated that the best prediction of the DA_PC_ model was the reinstatement behavior, although other behavioral outcomes could also be explained (Figure S7). These results align with the “dopaminergic hypothesis” of cocaine relapse and prior ideas proposing dopaminergic signaling in the ventral striatum as a major hallmark of drug-induced adaptations and a robust predictor of future relapse behavior ^37–39^. Moreover, we expand upon previous evidence by showing that low-dimensional features of the dopamine response space, pooling phasic responses from the first contact with the drug up until the reinstatement test, are *sufficient* to explain cocaine-seeking behavior in a relapse test.

### Sex refines the prediction of reinstatement behavior by multivariate patterns of dopamine release

*In vivo* microdialysis studies suggest that females exhibit higher cocaine-induced levels of extracellular dopamine in the dorsolateral striatum, but not in the NAc^40^. At baseline conditions, electrically-evoked phasic dopamine release is similar between sexes, although it is influenced by the estrous cycle^41^. To date, whether cocaine-evoked phasic dopamine responses evolve differently through the stages of the addiction cycle remains an open question of paramount relevance^42^. To explore this possibility, we examined whether sex played a role in dopamine-based predictions. If cocaine-evoked phasic dopamine responses – which are key to explain future relapse behavior – are influenced by sex, then considering this biological variable should refine the prediction of our DA_PC_ model. This was achieved by adding sex as an additional predictor, in combination with DA_PC1_, DA_PC2_ and DA_PC3_, to the MLR model of reinstatement behavior. Principal component scores were similar between males and females (Figure 3A), meaning that cocaine-evoked phasic dopamine responses did not diverge by sex when considering the animal’s history of cocaine self-administration as a whole. Despite this, our analyses indicated that sex contributed (sex; β = 0.65, *p <* 0.001) and improved (R^2^ = 0.82 > 0.73), the DA_PC_ model’s prediction of relapse behavior (Figure 3B, C). Likelihood ratio testing indicated that the model composed of sex and the DA_PC_ variables was preferred over the DA_PC_-only nested model (χ^2^ = 90.33, *p <* 0.001). According to AIC, the sex and DA_PC_ model represented a more parsimonious alternative, despite the additional free parameter (ΔAIC = −82.9; *p* < 0.001) (Figure 3D). These results reveal a previously unreported impact of sex as a predictor of cocaine-evoked dopamine dynamics and drug-motivated behavior. Our modelling approach indicates that sex interacts with individual differences in phasic dopamine responses to orchestrate future cocaine-seeking behavior during relapse.

**Figure 3.**
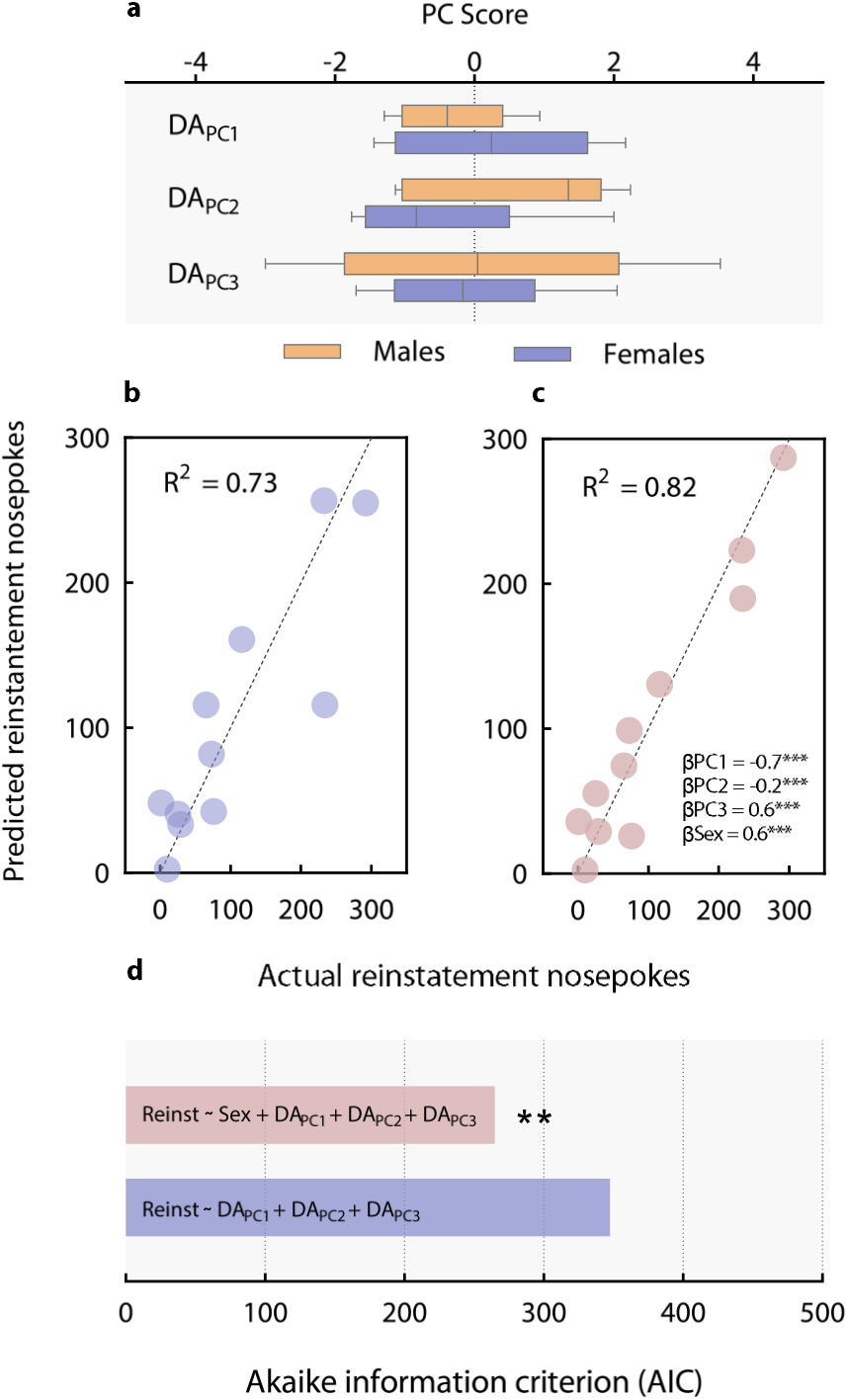
Inclusion of sex optimizes the prediction of reinstatement obtained by the DA_PC_ model. **a**) DA_PC1_,DA_PC2_, and DA_PC3_ principal component scores by sex (line = median, box = 95% CI, whiskers = min to max values). **b**) Actual vs. predicted reinstatement nosepokes obtained from a MLR model with DA_PC1_,DA_PC2_, and DA_PC3_ as covariates. The model’s R^2^ is shown as an inset. **c**) Actual vs. predicted reinstatement nosepokes obtained from MLR with sex, DA_PC1_,DA_PC2_, and DA_PC3_ as covariates. The model’s R and β-parameter estimates for sex and each DA_PC_ are shown as an inset. Asterisks indicate β significantly different from 0 (\*\*\**p* < 0.0001). **d**) Based on the Akaike information criteria, the sex + DA_PC_ model is less likely to be excluded. Asterisks indicate significant likelihood ratio test (\*\**p* < 0.01) favoring the sex + DA_PC_ model.

### Discrete phasic dopamine dynamics and latency to extinction are influenced by sex

Low-dimensional features of cocaine-evoked phasic dopamine responses provide information about future reinstatement behavior, especially when the subject’s sex is considered. In support of this evidence, we explored which discrete sex differences could be found in terms of behavior and cocaine-evoked dopamine events. Figure 4 illustrates averaged nosepokes, infusions, and related GrabDA_2m_ trial traces during the acquisition, maintenance, PR, extinction, and reinstatement phases of the self-administration experiment. Cocaine-evoked GrabDA_2m_ signals were compared using waveform (bootstrapped 95% CI procedure) and summary (averaged Δ F/F_0_ z-scores within time window of interest) analyses. Results show that both sexes consumed and pursued cocaine at similar levels (Figure 4A, B), as previously documented^28^. Despite a common behavioral phenotype, males exhibited higher dopamine release at cue onset and cocaine infusion (Figure 4C-D) than females (FR1). Behavioral differences were not seen during FR3 (Figure 4F, G), but males continued to show higher cocaine-evoked GrabDA_2m_ transients during cue presentation (Figure 4H-J). Waveform analyses showed post-infusion sex differences in dopamine release (Figure 4H), but summary analysis did not confirm this change (Figure 4J). The effect of sex during PR was diametrically opposed. Again, no behavioral differences were evident (Figure 4K, L), but reward-evoked accumbal dopamine release was higher in females (Figure 4M-O). However, there were no sex differences in terms of dopamine release at the time of cue onset (Figure 4N). Due to the constant violation of expectation and the increase in effortful demand that characterizes PR, within-session analyses can reveal information about dynamic changes in the dopaminergic encoding of rewards and reward-predictive cues. Thus, we focused our analyses on the early, mid, and late trials of PR (Figure 4P). Two-way repeated measures (RM) ANOVAs of cue-and cocaine-evoked dopamine release, with *sex* as the between-subject factor and *trial third* as the within-subject factor, resulted in no significant interaction effects (Figure 4Q, R). Therefore, we discarded sex differences in how dopamine release adapts to exponential changes in effort demand and prediction errors during operant cocaine seeking. Finally, we explored discrete sex effects during extinction and reinstatement of the cocaine-seeking behavior in the absence of the drug. There were no differences in terms of nose-poking behavior during the first extinction session (Figure 4S). Nonetheless, waveform analysis (bootstrapped 95% CI procedure) detected two significant increases in accumbal dopamine release, centered immediately before and after each nosepoke, in males (Figure 4T). No behavioral or dopaminergic differences were seen during the last extinction session (Figure 4U, V). Notably, male mice needed more sessions to reach extinction criteria, therefore showing higher resistance to extinguish cocaine-seeking behavior (Figure 4W). Despite no discrete differences in the number of nosepokes performed during cue-induced reinstatement of cocaine-seeking behavior (Figure 4X), males similarly exhibited increased dopaminergic encoding of the previously drug-paired cue (Figures 4Y). Our data clearly demonstrates that a closer examination at phasic dopamine responses obtained during cocaine self-administration reveals discrete sex differences. Specifically, we found that males showed greater accumbal dopamine transients upon cue presentation in virtually every experimental phase, regardless of whether cocaine was present or not. Interestingly, these dopaminergic differences existed in absence of a behavioral effect, with the exemption of the latency to extinguish cocaine seeking.

**Figure 4.**
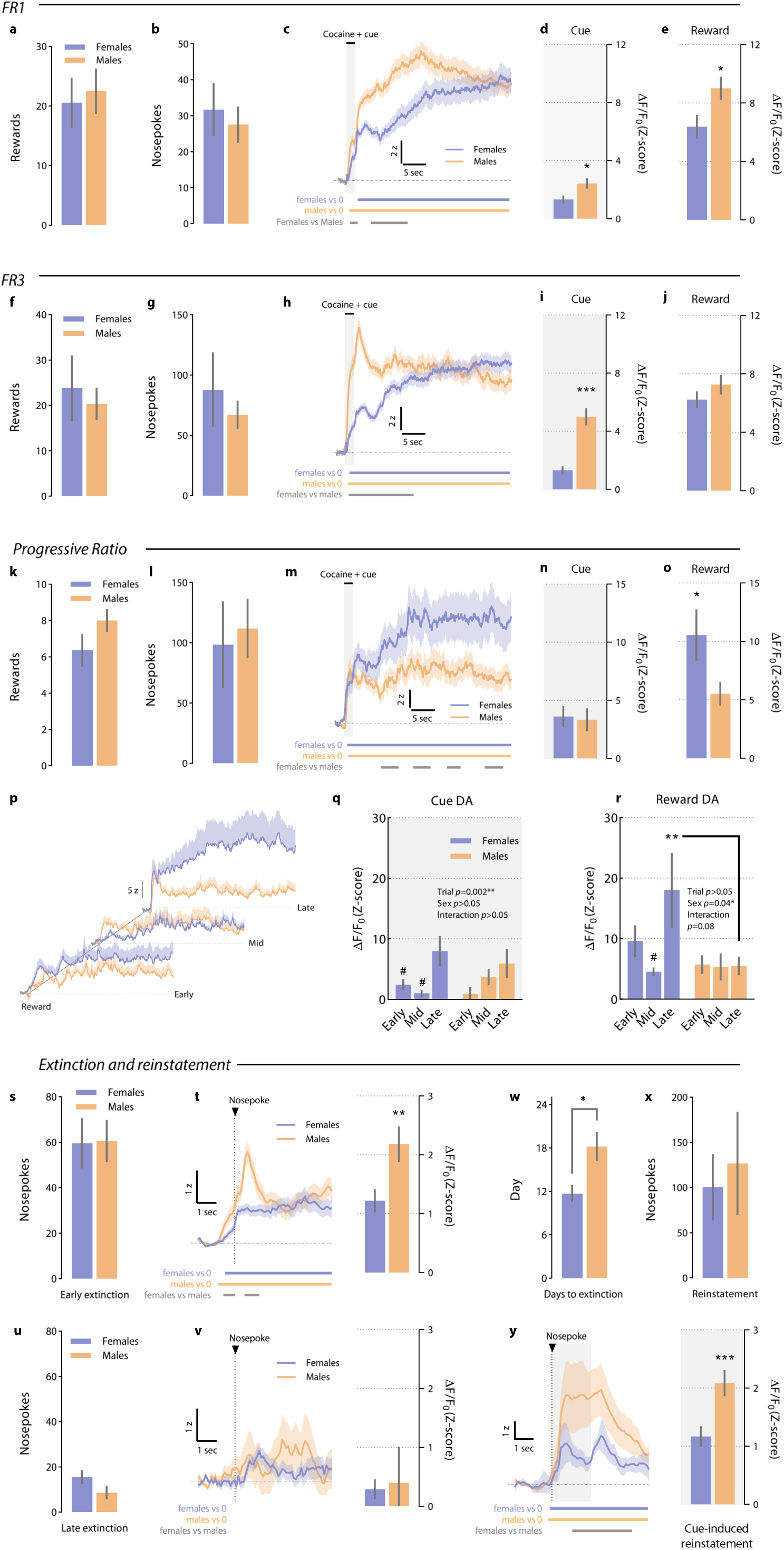
Discrete sex differences in cocaine-evoked phasic dopamine responses. **a, b**) Number of rewards obtained (*t*_12_ = 0.31, *p* = 0.76) and nosepokes given (*t*_12_ = 0.39, *p* = 0.70) segregated by sex during the early acquisition phase (FR1) (mean ± SEM active). **c**) NAc GrabDA_2m_ transients (mean ± SEM) segregated by sex centered −2 s to +30 s around cue onset and obtained during FR1 sessions. Colored bars below traces represent periods significantly different from 0 or between sexes, as defined by bootstrapped 95% confidence intervals (CIs). **d**) Mean ± SEM GrabDA_2m_ Δ F/F_0_ z-scores (0-2 s, cue onset) by sex from FR1 trials (*t*_277_ = 2.48, *p* = 0.013). **e**) Mean ± SEM GrabDA_2m_ Δ F/F_0_ z-scores (2-30 s, cocaine delivery) by sex from FR1 trials (*t*_278_ = 2.32, *p* = 0.021). **f, g**) Number of rewards obtained (*t*_12_ = 0.34, *p* = 0.73) and nosepokes emitted (*t*_12_ = 0.48, *p* = 0.63) by sex during the early acquisition phase (FR1) (mean ± SEM active). **h**) NAc GrabDA_2m_ transients (mean ± SEM) by sex centered −2 s to +30 s around cue onset and obtained during FR3 sessions. Colored bars below traces represent periods significantly different from 0 or between sexes, as defined by bootstrapped 95% CIs. **i**) Mean ± SEM GrabDA_2m_ Δ F/F_0_ z-scores (0-2 s, cue onset) by sex from FR3 trials (*t*_239_ = 6.08, *p* < 0.001). **j**) Mean ± SEM GrabDA_2m_ Δ F/F_0_ z-scores (2-30 s, cocaine delivery) by sex from FR3 trials (*t*_239_ = 1.17, *p* = 0.240). **k, l**) Number of rewards obtained (*t*_12_ = 0.61, *p* = 0.54) and nosepokes emitted (*t*_12_ = 0.12, *p* = 0.90) by sex during the progressive ratio (PR) test (mean ± SEM active). **m**) NAc GrabDA_2m_ transients (mean ± SEM) by sex centered −2 s to +30 s around cue onset and obtained during PR. Colored bars below traces represent periods significantly different from 0 or between sexes, as defined by bootstrapped 95% CIs. **n**) Mean ± SEM GrabDA_2m_ Δ F/F_0_ z-scores (0-2 s, cue onset) by sex from progressive ratio trials (*t*_76_ = 0.21, *p* = 0.83). **o**) Mean ± SEM GrabDA_2m_ Δ F/F_0_ z-scores (2-30 s, cocaine delivery) by sex from FR3 trials (*t*_75_ = 2.06, *p* = 0.042). **p**) Sex-specific NAc GrabDA_2m_ transients (mean ± SEM) from early, mid and late progressive ratio trials centered −2 s to +30 s around cue onset. **q**) Mean ± SEM GrabDA_2m_ Δ F/F_0_ z-scores (0-2 s, cue onset) by sex from early, mid and late PR trials (2-way RM ANOVA; *sex* (between-subjects), F_1,30_ = 0.07, *p* = 0.78; *trial* (within-subjects), F_2,41_ = 6.82, *p* = 0.002; *sex* x *trial*, F_2,41_ = 1.44, *p* = 0.24). Post-hoc comparisons: Bonferroni, # *p* > 0.05 vs. late trials, within same sex. **r**) Mean ± SEM GrabDA_2m_ Δ F/F_0_ z-scores (2-30 s, cocaine delivery) by sex from early, mid and late progressive ratio trials (2-way RM ANOVA; *sex* (between-subjects), F_1,30_ = 4.54, *p* = 0.041; *trial* (within-subjects), F_2,41_ = 2.63, *p* = 0.08; *sex* x *trial*, F_2,41_ = 2.59, *p* = 0.08). Post-hoc comparisons: Bonferroni, # *p* > 0.05 vs. late trials, within same sex; Bonferroni, ** *p* = 0.004 vs. males, late trials. **s**) Mean ± SEM nosepokes by sex (*t*_12_ = 0.06, *p* = 0.95) during the first extinction session. **t**) *Left panel*: NAc GrabDA_2m_ transients (mean ± SEM) by sex centered −2 s to +5 s around nosepoke and obtained during the first extinction session. Colored bars below traces represent periods significantly different from 0 or between sexes, as defined by bootstrapped 95% CIs. *Right panel*: Mean ± SEM GrabDA_2m_ Δ F/F_0_ z-scores (0-5 s, nosepoke) by sex from first extinction session trials (*t*_642_ = 2.75, *p* = 0.006). **u**) Mean ± SEM nosepokes by sex (*t*_12_ = 1.50, *p* = 0.15) during the last extinction session. **v**) *Left panel*: NAc GrabDA_2m_ transients (mean ± SEM) by sex centered −2 s to +5 s around nosepoke and obtained during the last extinction session. *Right panel*: Mean ± SEM GrabDA_2m_ Δ F/F_0_ z-scores (0-5 s, nosepoke) by sex from last extinction session trials (*t*_135_ = 0.21, *p* = 0.83). **w**) Mean ± SEM days required to reach extinction criteria by sex (*t*_12_ = 2.98, *p* = 0.011). **x**) Mean ± SEM nosepokes by sex (*t*_12_ = 0.40, *p* = 0.69) during the cue-induced reinstatement test. **y**) *Left panel*: Nac GrabDA_2m_ transients (mean ± SEM) by sex centered −2 s to +5 s around cue onset and obtained during the cue-induced reinstatement test. Colored bars below traces represent periods significantly different from 0 or between sexes, as defined by bootstrapped 95% CIs. *Right panel*: Mean ± SEM GrabDA_2m_ Δ F/F_0_ z-scores (0-2 s, cue onset) by sex from reinstatement trials (*t*_1121_ = 3.01, *p* = 0.002).

### Sex and patterned dopamine release predict the transition to extinction

Owing to the interactions between sex and dopamine signaling revealed previously, we next sought to reproduce the slower transition to extinction observed in males with the low-dimensional dopamine signatures DA_PC1_, DA_PC2_ and DA_PC3_. First, the sex-dependent effect on the latency to reach extinction criteria was confirmed by survival analyses. The proportion of females reaching extinction criteria across sessions was significantly higher at earlier timepoints than that of males (Logrank Mantel-Cox test; χ^2^ = 5.25, *p* = 0.021) (Figure 5A). Then we employed Cox proportional hazards regression (Methods) to model the transition to extinguished cocaine-seeking behavior across days. Sex and DA_PC1_, DA_PC2_ and DA_PC3_ were the main predictors. The Cox model accurately fit the observed extinction survival curves and recapitulated the slower extinction rates displayed by males (Figure 5B). Similarly, likelihood ratio testing indicated that the Cox model composed by sex and the DA_PC_ estimators explained the transition to extinction significantly better than an empty model with no covariates (χ^2^ = 12.42, *p* = 0.014), or an equivalent model without sex as predictor (χ^2^ = 9.65, *p* = 0.002). As a goodness-of-fit indicator, we used Harrell’s *C* statistic, which ranks how well the model attributed higher hazard ratios (risk for extinction) to subjects with shorter observation times (early transition to extinction). Harrell’s *C* ranges from 0.5 to 1, and scores over 0.7 are usually attributed to strong models^43^. The Cox regression model, with sex and the DA_PC_ predictors, yielded a strong prediction of the transition to extinction of cocaine-seeking behavior (*C* = 0.88). Hazard ratios were examined to determine the weight of each parameter in the model’s prediction of the extinction survival curve. In Cox proportional hazards regression models, hazard ratios (HR) are exponentiated versions of β-parameter estimates and characterize how much a predictor contributes to explain the probability of event transition when all other predictor variable values are held constant. Based on this metric, the Cox model indicated that sex (HR = 33.01) and DA_PC1_ (HR = 2.39), but not DA_PC2_ (HR = 0.88) or DA_PC3_ (HR = 0.58), contributed to explain the transition to extinguished cocaine-seeking. Of note, DA_PC1_ gathers GrabDA_2m_ fluorescence values from the early and late extinction sessions, wherein differences between sexes were clear. The above results reveal sex-specific trajectories of drug-seeking behaviors when access to the drug is withdrawn. Moreover, we document the importance of sex-specific phasic dopamine signatures that are sufficient to accurately predict the number of sessions needed to cease cocaine-seeking behavior.

**Figure 5.**
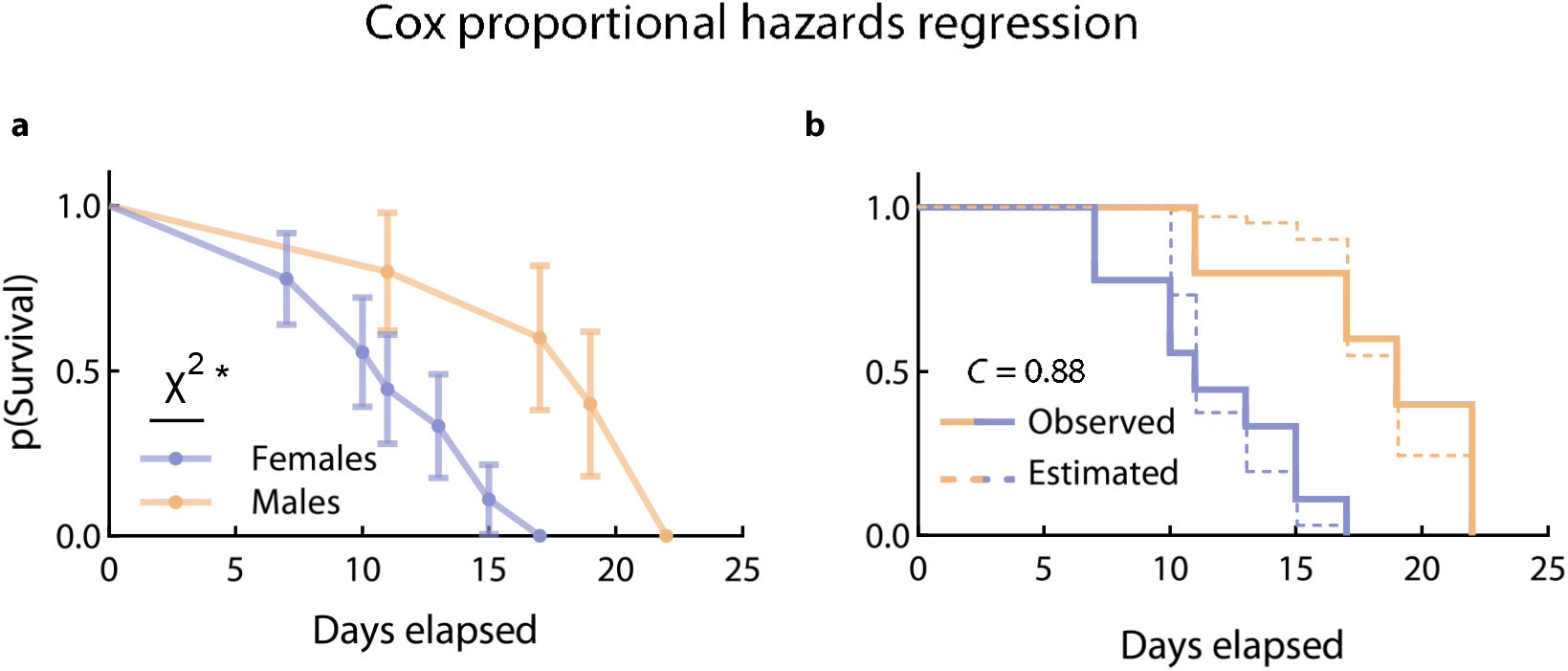
A combination of sex and dopamine responses are sufficient to predict resistance to extinction. **a**) Kaplan-Meier survival extinction curve of cocaine-seeking by sex (probability to not reach extinction criteria ± SEM) (Logrank Mantel-Cox test; χ^2^ = 5.25, \**p* = 0.021). **b**) Cox proportional hazard regression model composed by sex and DA_PC1_,DA_PC2_, and DA_PC3_ as covariates. Solid lines represent the observed Kaplan-Meier survival curves of extinction by sex. Dashed lines represent the estimated Kaplan-Meier survival curves by sex predicted by the Cox sex + DA_PC1_,DA_PC2_, and DA_PC3_ model. Harrel’s *C* statistic of goodness-of-fit is shown as an inset.

## Discussion

Cocaine hijacks dopaminergic signaling through dopamine transporter (DAT) blockade^44^, triggering long-lasting neuronal maladaptations^45^ that curtail control of instrumental behavior^46^. Altered dopaminergic signaling within the NAc has been extensively implicated in the progression to uncontrolled drug use and risk of relapse^47^. Prior work using electrochemical techniques showed that phasic dopamine release in the NAc encodes cocaine seeking^48–50^. More recently, subsecond dopamine transients aligned to cocaine intake were documented using a genetically-encoded fluorescent reporter^51^. However, surprisingly little is known about the dynamics of drug-evoked phasic dopamine release throughout the different stages of drug intake and addiction^52^. Here, we take advantage of novel photometric techniques to parse out dopamine dynamics during cocaine seeking throughout the animal’s entire experience with the drug, to expand our knowledge of the relationship between this monoamine and relapse.

We report, for the first time, stable recordings of phasic dopamine activity, from acquisition to the reinstatement of cocaine-seeking behavior in freely moving mice. The long-term, concurrent use of fiber photometry and i.v. drug self-administration in C57BL/6 mice opens the possibility to implement genetic manipulations in longitudinal addiction studies. It is worth noting that prior studies conducted similar long-term NAc recordings but were exclusively focused on escalation of intake^53,54^, based on a within-session extinction procedure^55,56^, or during craving incubation^57^. Despite key methodological differences (species, cocaine doses, and delivery times of drug infusion and cue presentation), NAc dopamine transients documented here are equivalent to those previously reported^31,53–55^, especially when using an identical GrabDA_2m_ sensor^51^. Cocaine-evoked increases of NAc dopamine occupation of the sensor (assessed as fluorescence) were of longer duration compared to a natural reinforcer (sucrose).

Long-term GrabDA_2m_ recordings were surprisingly effective at predicting nose-poking behavior during the reinstatement test after a period of extinction training. Persistent risk of relapse is a defining feature of drug addiction, and, despite relevant alternatives, it remains a primary endpoint for treatment of substance use disorders^58,59^. Thus, we tested whether we could predict each animal’s reinstatement behavior by their dopaminergic signatures. Our results indicate that future relapse behavior was correlated with the amplitude of cocaine evoked NAc dopamine transients observed on the first FR1 sessions (acquisition). This suggests that mesolimbic dopamine responses during early access to cocaine (intoxication/appetitive phase) carry information about future behavioral risk to relapse. More importantly, we uncovered a much stronger correlation between cue-evoked dopamine release observed during PR responding and relapse. This evidence supports early work postulating that PR schedules of reinforcement are particularly well-suited to infer maladaptive cue encoding of drug reinforcement^34,35,60^. Owing to the increasing availability of positron emission tomography (PET) tools^61^, such a relationship could avail PET estimations of NAc dopamine markers in a PR schedule as powerful indicators of future relapse in concurrent cocaine users. Building on this finding, we next sought to optimize the information gathered by our GrabDA_2m_ longitudinal measures of dopamine dynamics to obtain a comprehensive linear model of relapse. Using a PCR approach, we showed that patterned NAc dopamine release explained cue-induced reinstatement behavior to a high degree of accuracy. Indeed, these dopamine signatures were more informative than a similar MLR model using the animal’s behavior as predictor, consistent with the notion that reinstatement to cocaine-seeking behavior is associated with NAc dopamine signaling and the maladaptive plasticity that accompanies it^39^. Our results further support the sufficiency of drug-influenced phasic dopamine dynamics for enhanced risk of relapse. Using this approach, we effectively expanded the space of variables by assessing the history of dopamine responses throughout the entire self-administration experiment. In doing so, we report the importance of early dopaminergic responses associated with cocaine and expand our comprehension of the monoaminergic mechanisms leading to cocaine relapse^62^.

Trajectories of drug use and abuse are markedly influenced by gender and sex^63^. In 2020, of the 68,830 opioid-related overdose deaths registered in the US, 70% were among men^64^. Despite lower death rates, women transition faster from recreational drug use to problematic drug abuse and exhibit higher risk of relapse^65^. Preclinical studies have addressed these disparities in laboratory animals, and the prevailing notion is that female rodents show increased drug use escalation and facilitated relapse^42^. However, a recent exhaustive review concluded that there is not enough evidence supporting the idea that women and female rodents are more vulnerable to cocaine craving and relapse^23^. Here, we tested the DA_PC_ model of relapse behavior to investigate possible sex differences present in our dataset. In support of prior reports^66–70^ (but see^71^), we could not find evidence in favor of increased vulnerability to cue-induced reinstatement of cocaine-seeking in female mice. Indeed, males displayed increased dopamine fluorescence on discrete phases of voluntary seeking and intake of cocaine more readily (acquisition, maintenance, early extinction, and reinstatement). Moreover, other studies have suggested greater accumbal dopaminergic responses to cocaine and d-amphetamine in males, regardless of hormone treatment^72,73^. This is an important detail because during the estrous phase of the menstrual cycle, or hormonal treatment (e.g., estradiol), female rodents showed potentiation of mesoaccumbal dopamine neuron activity and cocaine’s rewarding effects in a conditioned place preference study^41^. In our dataset, females displayed greater dopamine release during the PR task. Although we did not assess the menstrual cycle, these results reveal evolving sex differences in dopaminergic encoding of cocaine seeking. Nevertheless, low-dimensional features of dopamine responses (DA_PC1_, DA_PC2_ and DA_PC3_) were equivalent between sexes. The co-occurrence of discrete dopamine differences with a common cocaine-evoked dopamine response trait (DA_PC1,2,3_) may seem contradictory, but it is reasonable to think that both sexes followed trajectories that are different, but comparable as a whole, in order to ensure a common behavioral adaptation to a given environmental setting. Thus, sex does not determine a divergent behavioral adaptation between males and females, but rather, acts in concert with other individual differences to leverage a shared behavioral outcome. Here, we show that when sex is incorporated into the DA_PC_ regression model, an improvement of the predictive performance is observed. Notably, the combination of sex and the patterned dopamine release yielded a remarkably accurate prediction of reinstatement nosepokes, reproducing 82% of the observed variance with a relatively simple linear regression that did not consider many dopaminergic algorithms^62,74^ associated with motivated behavior.

We also observed a greater resistance to cease responding under extinction conditions in males. The literature addressing sex-dependent extinction trajectories is still scarce, but some reports previously point to opposite changes^70^ or no differences between sexes^75^. Unlike these studies, we characterized slower extinction rates by using survival analyses, which relied on each animal’s baseline responding (dependent on extinction criteria) and thus yielded a normalized measurement. It is important to keep in mind that there is a considerable conceptual distance between the extinction procedure followed here and the abstinence period typically preceding relapse episodes in humans^76^. Nevertheless, investigating the behavioral and neural mechanisms of extinction can increase our understanding of drug seeking reduction^77^. The resistance to reduce cocaine seeking by male mice aligns well with human reports suggesting a higher propensity in men to transition from voluntary abstinence to cocaine use (relapse)^25^. In addition, we showed that males’ slower extinction rates were accompanied by increased NAc dopamine release during cocaine seeking at the beginning of extinction training and during memory retrieval (reinstatement). Recent work revealed that activation of NAc cholinergic interneurons promotes extinction of cocaine-preference memories^78^. Considering that dopamine desynchronizes cholinergic interneuron activity^79^, we postulate that the resistance to extinction observed in males is related to their increased dopaminergic encoding and the resulting disruption of NAc cholinergic interneuron mediation of extinction learning. Importantly, sex and patterned dopamine responses accurately predict the probability to extinguish cocaine seeking. Thus, our results reveal a previously undescribed predictive power of patterned dopamine responses (DA_PC1_) at drug-associated cues, sampled throughout a history of cocaine self-administration, on the progression to reinstatement. Finally, it is worth noting that this finding provides evidence for male vulnerability to reinstate cocaine-seeking behavior upon drug-paired cue re-exposure.

In conclusion, our results greatly expand and enrich our knowledge of the role of NAc dopamine signaling in relapse to cocaine seeking. Our experiments unambiguously show that it is possible to recapitulate reinstatement and extinction with a simple linear model that integrates the animal’s longitudinal repertoire of cocaine-evoked phasic dopamine responses in the NAc. A closer examination of the DA_PC_ model reveals the particular importance of early dopamine responses (maintenance, PR) in the gating of future relapse behavior. Fiber photometry in self-administering mice allowed us, for the first time, to characterize sex-specific phasic dopamine responses during the pursuit of cocaine. We postulate that this validated neuromarker, now accessible through human PET studies, could be used to refine patient prognosis in both research and clinical settings.

## Methods

### Animals

For all the experiments, female and male C57BL/6J mice were used (N = 12). Prior to fiber implantation surgery, mice were group-housed in plastic cages with ad libitum access to food and water. Following surgery, animals were singly-housed. All rodent holding rooms were maintained at 24 °C and 40–50% humidity under a 12-h light/dark cycle with lights on at 07:00 h. Research facilities were certified by the Association for the Assessment and Accreditation of Laboratory Animal Care (AAALAC), and experimental procedures were approved by the Institutional Animal Care and Use Committee (IACUC) at University of Maryland School of Medicine and in accordance with the Guide for the Care and Use of Laboratory Animals^80^. Animals were transduced with a genetically-encoded fluorescent dopamine sensor^17^ (GrabDA_2m_) and implanted with optical fibers in the core part of the NAc on postnatal day (PND) 70. pAAV-hsyn-GRAB_DA2m was a gift from Yulong Li (Addgene viral prep #140553-AAV9). On PND 100, they underwent the second surgery procedure, in which they were implanted with intravenous catheters and trained to self-administer cocaine. Then, mice were simultaneously recorded for *in vivo* fiber photometry and cocaine self-administration.

### Viral delivery and optical fiber implantation

Animals were anesthetized using isoflurane in O_2_ (4% induction and 2% maintenance) and then placed on the stereotaxic apparatus. To express fluorescent dopamine-sensor (GrabDA_2m_) in the NAc, AAV9-hSyn-DA-sensor 4.4 (10^13^ genome copies (gc)/ml)) was unilaterally injected using the following coordinates: antero-posterior, 1.1 mm; medio-lateral, 1 mm; and dorsal-ventral, −3.8 mm; relative to bregma. By a microsyringe pump, AAV9-hSyn-DA-sensor 4.4 (500 nl) was slowly infused thorough a sharp glass pipette into the target brain area at rate of 100 nl/min. Immediately following virus infusion, an optical fiber (diameter, 400 µm; NA, 0.5; Thorlabs) bounded to a ceramic ferrule was implanted with its tip targeting 0.1 mm above the abovementioned NAc DV coordinates. Ceramic ferrules were secured to the skull using dental cement with the help of two skull-penetrating screws. Mice were gently removed from the stereotaxic instrument and placed over a heat pad in their home cages. With daily monitoring for wound healing, mice were singly housed and allowed to recover for 4 weeks.

### Intravenous catheterization surgery

After 4 weeks of recovery and viral expression, GrabDA_2m_ fluorescence was tested during environmental exploration. Only those animals showing fluctuation in fluorescence intensity were selected for catheterization of jugular vein. Surgical implantation of the catheter into the jugular vein was performed following anaesthetization with a mixture of Ketamine hydrochloride (100 mg/kg) and Xylazine hydrochloride (10 mg/kg), injected in a volume of 0.1 mL/10 g body weight, i.p. Indwelling i.v. silastic catheters (0.3 mm inner diameter, 0.6 mm outer diameter) were implanted 1.3 cm into the right jugular vein and anchored with suture^81^. The remaining tubing ran subcutaneously to the cannula, which exited at the midscapular region. All incisions were sutured and coated with antibiotic ointment (Bactroban, GlaxoSmithKline). After surgery, animals were allowed to recover for 3 days prior to initiation of self-administration sessions. To maintain patency, catheters were flushed daily with heparinized saline (30 USP units/ml).

### Intravenous cocaine self-administration

After surgery recovery, mice were trained in operant chambers (Model ENV-307A-CT, Med Associates) equipped with two holes, one randomly selected as the active hole and the other as the inactive. Cocaine 0.5 mg/kg/ was delivered in a 20 μl injection over 2-s via a syringe mounted on a microinfusion pump (PHM-100A, Med-Associates), a single-channel liquid swivel (375/25, Instech Laboratories) and the mouse’s intravenous catheter. All FR1 and FR3 sessions started with a cocaine priming infusion. When mice responded on the active hole, the stimulus lights (located above the nosepoke hole) and tone were presented for 2-s and a cocaine infusion was delivered automatically over these 2-s. Each infusion was followed by a 10-s time-out period in which a nosepoke on the active hole had no consequences but was recorded.

#### Acquisition and maintenance of operant cocaine taking

Mice were trained to nosepoke in order to receive 0.5 mg/kg cocaine infusions under a fixed ratio 1 reinforcement schedule (FR1) in 2h-long daily sessions. No food self-administration pre-training was conducted. Mice were moved to an FR3 reinforcement schedule when the following criteria were met on 2 consecutive FR1 sessions: *a*) ≥ 65% of responses were received at the active hole; and *b*) a minimum of 15 responses on the active hole. After meeting criteria, animals underwent 5 more FR3 sessions.

#### Progressive ratio test session

After the five FR3 sessions, animals were tested in a PR schedule, wherein the response requirement to earn infusions escalated according to the following series: The PR session ended when mice were not able to earn the response requirement in 1-h and was performed only once.

#### Extinction and reinstatement of cocaine seeking

After PR testing, mice underwent operant extinction of cocaine seeking behaviors. During these sessions, nosepokes in the active hole produced neither cocaine infusion nor cue presentation. Extinction sessions (2-h long) were conducted once a day, 5 days/week until reaching the extinction criteria in two consecutive days (performing less than 15 responses or 40% of the mean nosepokes exhibited during FR3 testing). Twenty-four hours after, mice underwent a cue-induced reinstatement session, in which they were confined to the operant chambers for 2-h. During the reinstatement session, mice did not receive cocaine infusions but nose-poking in the active hole resulted in cue light presentation and activation of the cocaine microinfusion pump under a FR1 schedule.

### Fiber photometry recording

Fiber photometry of GrabDA_2m_ signals was conducted in all cocaine self-administration sessions, from acquisition to reinstatement. Two fiber-coupled LEDs producing 470 nm (M470F3, ThorLabs) and 405 nm (M405FP1, ThorLabs) lasers were used as the excitation source. The laser beams were reflected and coupled into a fluorescence minicube (FMC4, Doric Lenses). A 2 m-long optical fiber (400 µm, Doric Lenses) was used to transmit light between the fluorescence minicube and the implanted fiber. The laser intensity was measured at the tip of optical fiber and adjusted to ∼ 5 µW and ∼10 µW for the 405-nm and 470-nm lasers, respectively. GrabDA_2m_ fluorescence was collected by the optical fiber, passed through the fluorescence minicube, and projected onto a photoreceiver (Newport 2151, Doric) where light intensity was converted into current signal. A RZ5P real-time processor (Tucker-Davis Technologies, TDT) was used to convert the current signal to voltage signal, which was processed through a low-pass filter (6 Hz, 6^th^ order Butterworth filter) to allow filtering of noise at higher frequency. Finally, voltage signal and transistor–transistor logic (TTL) signals coming from the operant chambers were sampled at 330 Hz and recorded using the Synapse software (ThorLabs). Cue presentation onset, cocaine infusion offset and nosepoke onset times in the active and inactive holes were recorded as independent TTLs.

### Fiber photometry signal processing and PETH creation

GrabDA_2m_ signals were processed using a custom-written MATLAB program, based on Barker et al.^82^ (available at www.tdt.com/support/matlab-sdk). Noise-related changes in fluorescence across the whole experimental session were removed by scaling the isosbestic control signal (405 nm) and regressing it onto the dopamine-sensitive signal (470 nm). This regression generated a predicted model of the noise that was based on the isosbestic control. Dopamine-independent waveforms on the 405 nm model were then subtracted from the raw GrabDA_2m_ signal to remove movement, photo-bleaching, and fiber-bending artifacts. PETHs were constructed using 100-ms bins surrounding the event of interest. PETH time windows for FR1, FR3 and PR recordings spanned −2s to 30s, centered around cue presentation TTLs. Time windows for extinction and reinstatement recordings spanned −2s to 5s, centered around nosepoke or cue presentation TTLs, respectively. Each bin of the PETH was z-scored by subtracting the mean fluorescence found in the −2s to −1s time window preceding each trial and dividing by the s.d. across those windows (n = number of trials). In the FR1, FR3, PR and reinstatement phases of the experiment, z-scores obtained during the first two seconds following cue presentation (0-2 s, aligned to cue onset) were considered to reflect cue-evoked dopamine-based fluorescence. When cocaine was available, z-scores of the remaining period (2-30 s, aligned to cocaine delivery offset) were interpreted as reward-evoked dopamine-based fluorescence. For the extinction recordings, the two seconds following each nosepoke (0-2 s) were interpreted to reflect nosepoke-related dopamine-based fluorescence. Significant dopamine transients while cocaine was available often exceeded the 30-s time window used on the PETHs. To visualize the full extent of the dopamine transients, we plotted a session-wide GrabDA_2m_ recording from a representative FR1 session with overlayed infusion timestamps (Figure S1). Figure S1 shows that cocaine-evoked GrabDA2m transients can last for several minutes before decaying to baseline. Inter-response intervals followed a similar temporal distribution, indicating that only a residual number of extra responses fell within the 30-s time window used for the PETH analyses. Delving further into this temporal coincidence, we modeled predicted quantities of cocaine brain concentrations (C_brain_) using the equation *A*(*e*^−*αt*^ − *e*^−*βt*^), as characterized by Pan et al.^33^, where A = 4.82 (a multiplicative factor including the cocaine dose), α = 0.01, β = 0.0095 and *t* being the time in minutes since the previous cocaine infusion. Except for the initial 5 minutes of the session, corresponding to the “loading phase” of consumption, cocaine-taking responses coincided with the decrease of cocaine content in the brain (C_brain_). *ΔF/F*_*0*_ values were expressed as z-scores.

### Statistical tests and GrabDA_2m_-based predictive models

We analyzed the results of single factor, two-group, parametric variables (GrabDA_2m_ z-scores and nosepoke comparisons between sexes) with unpaired Student’s *t*-tests. Parametric measures resulting from the combination of two factors (PR z-scores by sex) were analyzed with two-way ANOVAs. When an experimental condition followed a within-subject design (e.g., trial, session) an ANOVA with repeated measures was calculated. In addition to z-score summary analyses, GrabDA_2m_ fluorescence values were compared using waveform analyses, hence providing temporally-defined significance. Waveform analyses were based on the bootstrapping CI procedure developed by Jean-Richard-dit-Bressel et al.^29^. Significant transients within the PETH were defined as periods (with a minimum length of 0.5-s) whose bootstrapped 95% CI did not contain 0 (baseline) or the other group’s waveform. Bootstrapped CI were obtained by randomly re-shuffling (1000 boot-straps) trial z-scores. The bootstrap distribution was then expanded by a factor of 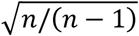 to adjust for narrowness bias^29^.

We then estimated Pearson’s correlation coefficients between dopaminergic and behavioral variables, with an emphasis on reinstatement incidence, to uncover initial relationships. These variables were defined as averaged number of nosepokes or averaged cue-, reward- or nosepoke-evoked z-scores obtained during FR1, FR3, PR, extinction and reinstatement recordings.

To generate a dopamine-based MLR model of reinstatement nosepokes, we followed a PCR strategy that allowed us to minimize the number of free parameters while pooling information from nine different dopaminergic variables. In addition, PCR enabled an unbiased approach in which no prior hypothesis was needed to cherry-pick the most interesting dopaminergic variables to fit into the linear regression. First, a low-dimensional representation of all the dopaminergic variables was obtained by PCA^83^ after scaling the data to have a mean of 0 and s.d. of 1. The computed principal components compiled multivariate patterns of NAc dopamine responses to cocaine throughout the animal’s entire history of drug exposure. The resulting loading scores are shown in Figure S4. Second, we selected the first three principal components (referred to as DA_PC1_, DA_PC2_ and DA_PC3_), which explained nearly 75% of the observed dopaminergic variance, to serve as covariates in a MLR model of reinstatement nosepokes. We replicated our main result with two other principal component selection methods^84^ (Figure S6). The “Kaiser rule” method selected four principal components with an eigenvalue higher than 1. The “Elbow rule” method selected the first five principal components before an apparent dip in the explanatory power of the next factor. Given that the number of nosepokes during reinstatement is a count of events (with no possible negative or decimal values) and that its distribution violated the normality assumption (tested with three different statistics; Shapiro-Wilk, W = 0.248, *p* = 0.048; Anderson-Darling, A2* = 0.712, *p* = 0.044 and Kolmogorov-Smirnov, KS distance = 0.218, *p* = 0.046) we fit the MLR model using a Poisson’s regression. The dopamine-based model (DA_PC_) of reinstatement behavior came thus defined by ln(*Ŷ*) = *β*_0_ + *β*_1_(*DA*_*PC*1_) + *β*_2_(*DA*_*PC*2_) + *β*3_3_(*DA*_*PC*3_). Similarly, we obtained the same Poison’s MLR model of reinstatement nosepokes using principal components derived from the behavioral variables shown in Figure 2B as covariates (BE_PC_). To compare goodness-of-fit between models we used the small-sample corrected AIC, which accounted for both the percentage of variance explained and the different number of free parameters included. The difference between AIC was calculated by *ΔAIC* = 2 ln(*LR*) + 2*Δdf*, where LR was the negative log-likelihood ratio value of the model and *df* the number of free parameters. The probability of the alternative model (BE_PC_) being more likely to be correct than the reference model (DA_PC_) was determined by *p* = 1 − (*e*^0.5Δ*AIC*^ /1 + *e*^0.5*ΔAIC*^). Nested models (e.g., DA_PC_ ± sex) were compared using both ∆AIC and likelihood ratio methods.

To replicate our MLR results, we performed an additional Bayesian Poisson inference procedure^85^ to predict reinstatement nosepokes using DA_PC1_, DA_PC2_ and DA_PC3_ as regression coefficients. First, we used Markov chain Monte Carlo (MCMC) method to generate 10000 samples (first 2500 iterations discarded as burn-in) that followed the posterior distributions of reinstatement nosepokes – the observed data. Then, we ran a Metropolis-Hastings algorithm that iteratively sampled the β coefficients from the simulated samples until getting with the β parameters that most closely regressed the observed data. The logarithm of the pseudo-marginal likelihood (LPML)^86^ and R^2^ between the observed and a randomly sampled distribution of reinstatement values were used as goodness-of-fit measures. Bayesian Poisson regression was performed under the “bpr” R package (Windows 11).

To characterize sex differences in the progression to extinguished cocaine-seeking behavior, Kaplan-Meyer survival curves were computed. Then, multivariate patterns of NAc dopamine release (DA_PC1_, DA_PC2_ and DA_PC3_), and sex, were used as covariates to recover the observed Kaplan-Meyer survival curves. This was achieved using a semi-parametric Cox proportional hazards regression, which defined the model 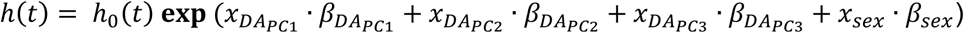, where *h*(*t*) is the estimated hazard at time *t*, h_0_(*t*) is the baseline hazard when all the predictors are equal to zero, *x*_i_ are the values of the predictor variables and β_i_ the parameter coefficients. Harrell’s *C* statistic, equivalent to the area under the ROC curve for logistic regressions, was used to define the goodness-of-fit for the Cox model of extinction^87^.

## Funding acknowledgements

This research was supported by the National Institute on Drug Abuse (R01 DA022340 and R01 DA042595 to JFC; F32 DA039690 to JMW; and F32 DA043967 and K99/R00 DA047419 to NEZ) and by the São Paulo Research Foundation, FAPESP (2019/23286-3; SAE). MAL received a FPU grant funding (15/02492) from the Ministerio de Educacion, Cultura y Deporte (Gobierno de España).

## Supplementary Figures

**Figure S1.**
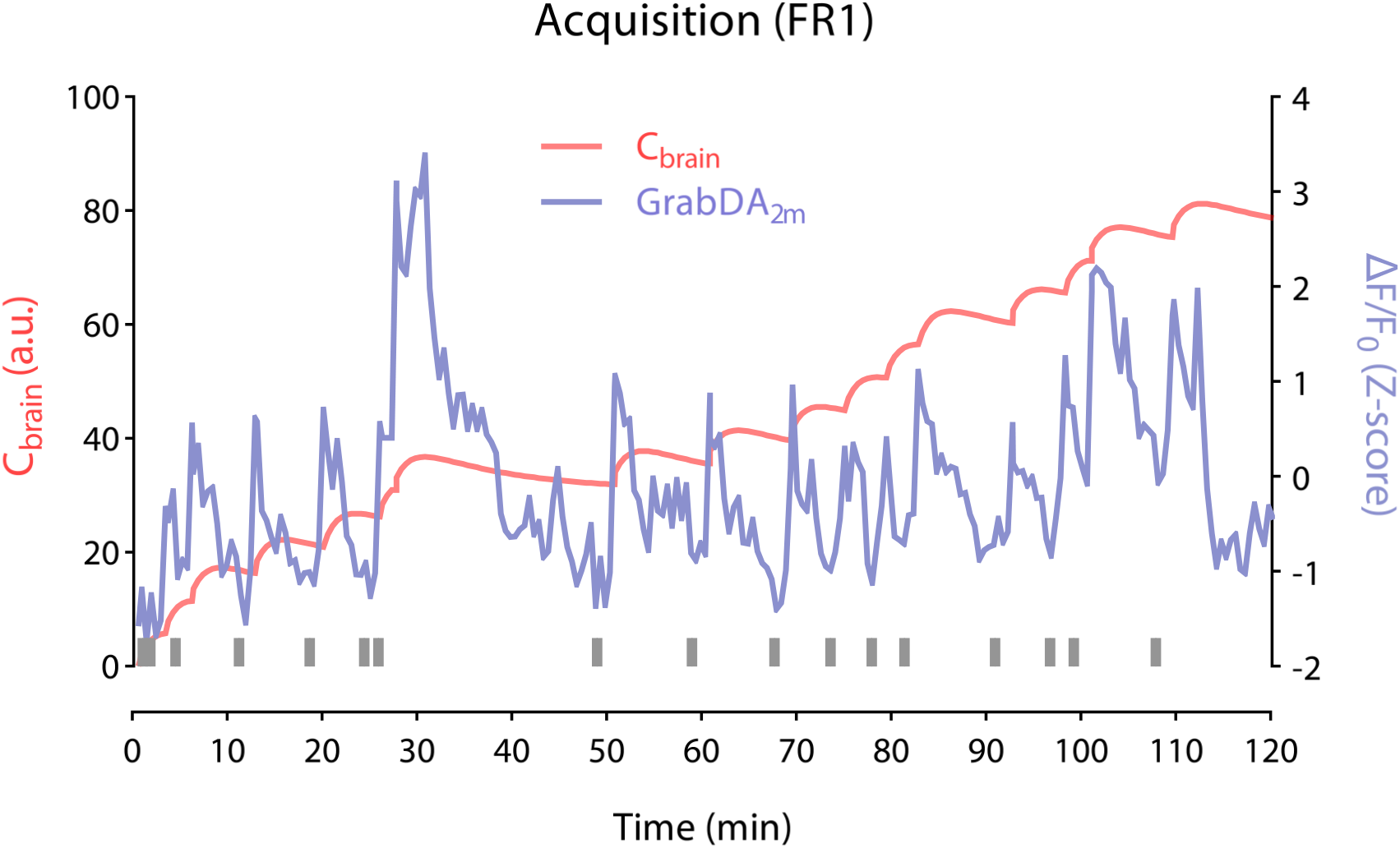
Representative GrabDA_2m_ fluorescence trace and modeled cocaine brain concentrations during a self-administration FR1 session. Bottom grey bars represent cocaine infusions timestamps.

**Figure S2.**
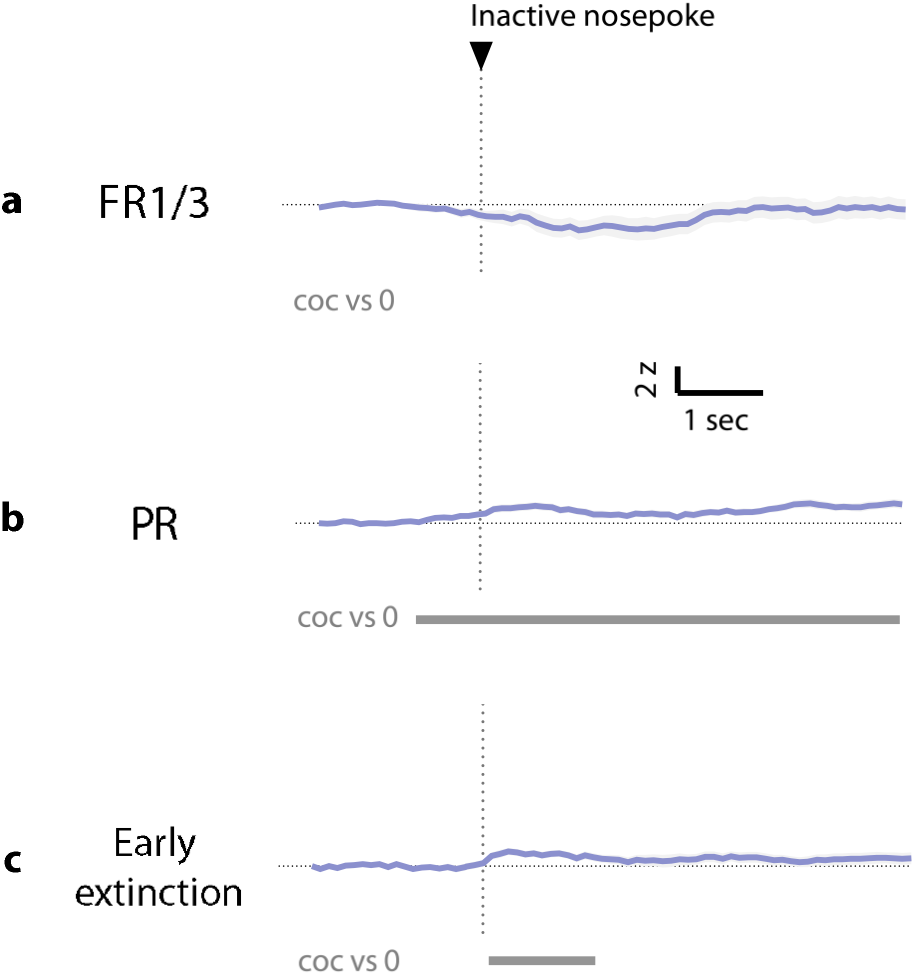
Dopaminergic encoding of inactive nose-poking responses in the NAc during the phases in which a pre-cue significant transients were detected. (a) Mean ± SEM GrabDA_2m_ transients (Δ F/F_0_ z-scores) aligned to inactive nosepokes during FR1 and FR3 (combined). (b) Mean ± SEM GrabDA_2m_ transients aligned to inactive nosepokes during PR. A small but significant transient is detected throughout the initiation of the motor sequence leading to the inactive nose-poke and the subsequent five seconds. (c) Mean ± SEM GrabDA_2m_ transients aligned to inactive nosepokes during early extinction. The pre-cue dopamine transient detected in active nosepokes is not present during inactive nosepokes of the same phase. Traces centered −2 s to +5 s around inactive nosepoke onset. Bottom grey bars below traces represent periods significantly different from 0, as defined by bootstrapped 95% confidence intervals.

**Figure S3.**
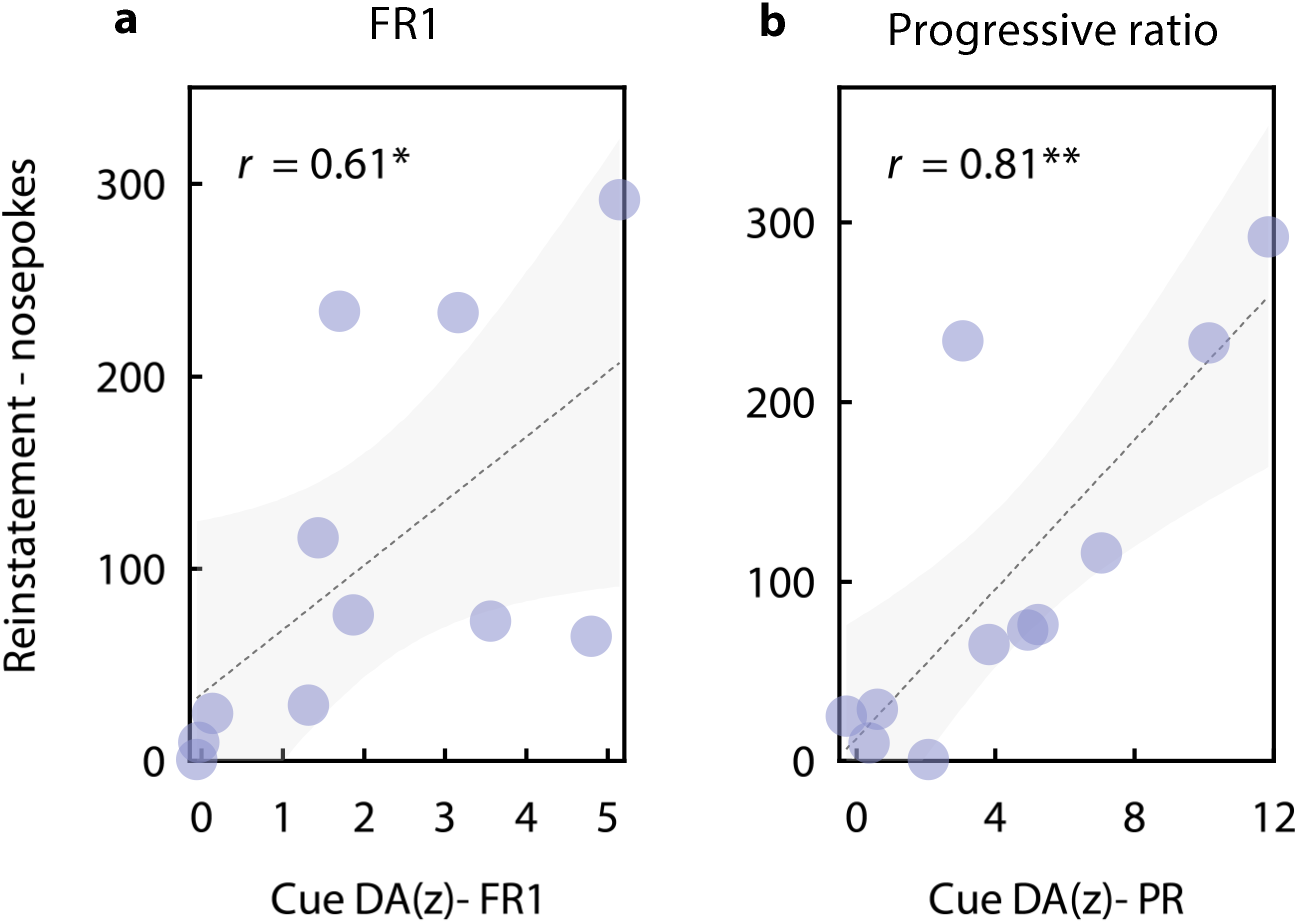
Significant Pearson’s coefficients between reinstatement incidence and dopamine-related cocaine responses. (a) Cue-evoked NAc GrabDA_2m_ Δ F/F_0_ z-scores (mean ± SEM) (0 – 2s, cue onset) from FR1 trials are positively correlated with the number of nosepokes given during the reinstatement test. (b) Cue-evoked NAc GrabDA_2m_ transients (mean ± SEM) (0 – 2s, cue onset) from PR trials are positively correlated with the number of nosepokes given during the reinstatement test. Shaded areas depict 95% CI of the regressed slope. Two-tailed Pearson’s r; *\*p* = 0.047; *\*\*p* = 0.003.

**Figure S4.**
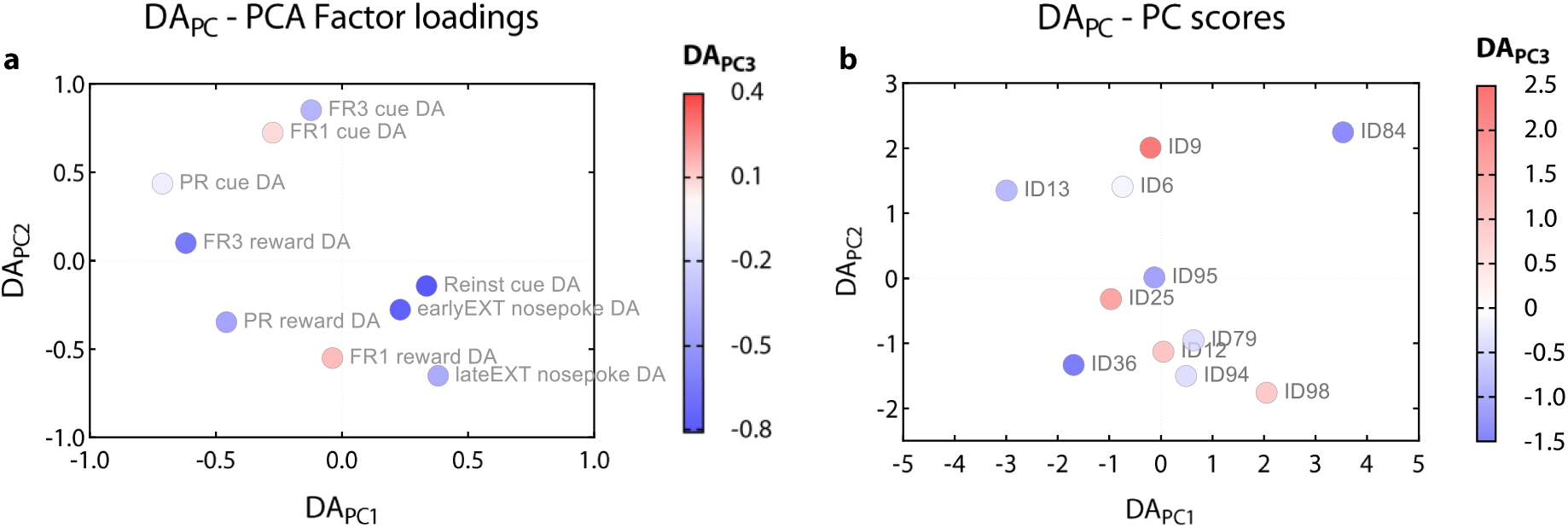
DA_PC_ factor loadings and subject’s scores. (a) Dopaminergic response variables obtained throughout cocaine self-administration contribute in different directions to the select principal components DA_PC1_, DA_PC2_, and DA_PC3_. Each variable’s empirical value (ΔF/F_0_ z-scores) contributes to each principal component score with the weighting factors depicted in Figure 2C-E. Variables with positive loadings for a given principal component increase the animal’s score on that factor (e.g., reinstatement cue-evoked dopamine release for DA_PC1_). Variables with negative loadings decrease the animal’s score on that factor (e.g., FR3 cue-evoked dopamine release for DA_PC1_). Factor loadings are crucial to understand the directionality of the contribution that each empirical variable has on the MLR model of reinstatement behavior considering the β-parameters of each DA_PC_ (shown in Figure 2G). (b) Principal component scores of each subject in the dopaminergic space obtained after dimensionality reduction.

**Figure S5.**
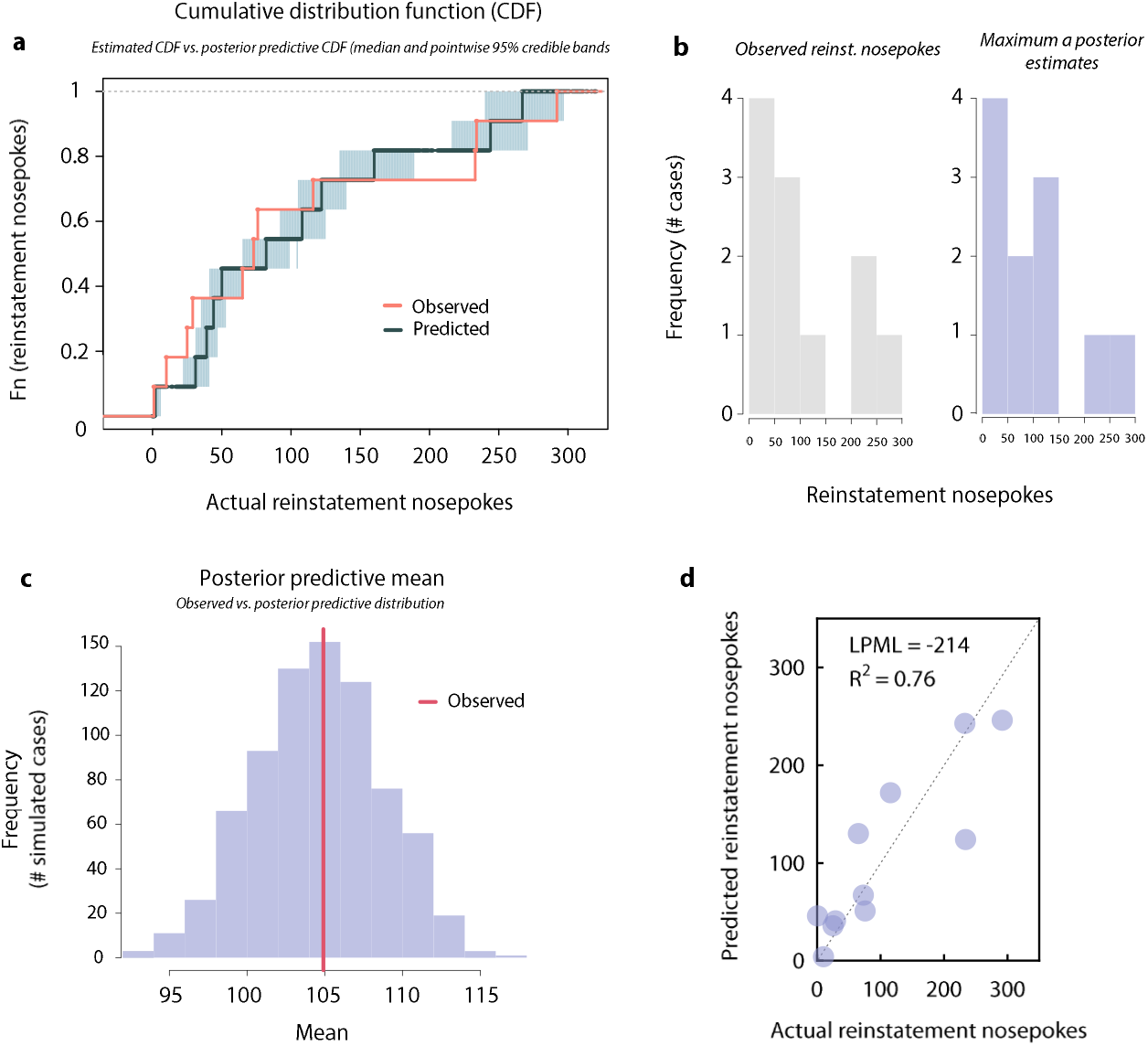
Bayesian Poisson inference of cued reinstatement behavior using multivariate patterns of NAc dopamine release as regression coefficients. **a**) Posterior predictive check comparing the empirical cumulative distribution function (ECDF) of the observed data and the cumulative distribution function computed on the simulated predictive samples. **b**) Reinstatement incidence distribution of the observed data and the maximum *a posteriori* predictive distribution. **c**) Comparison between the simulated mean distribution (among the simulated predictive samples) and observed mean of reinstatement nosepokes. **d**) Biplot of the observed reinstatement nosepoke values and a randomly selected simulation from the Metropolis-Hasting-sampled distribution. Log pseudo-marginal likelihood (LPML) and R^2^ are shown as metrics of goodness-of-fit. R^2^ obtained by regressing observed reinstatement values on the randomly-selected sample.

**Figure S6.**
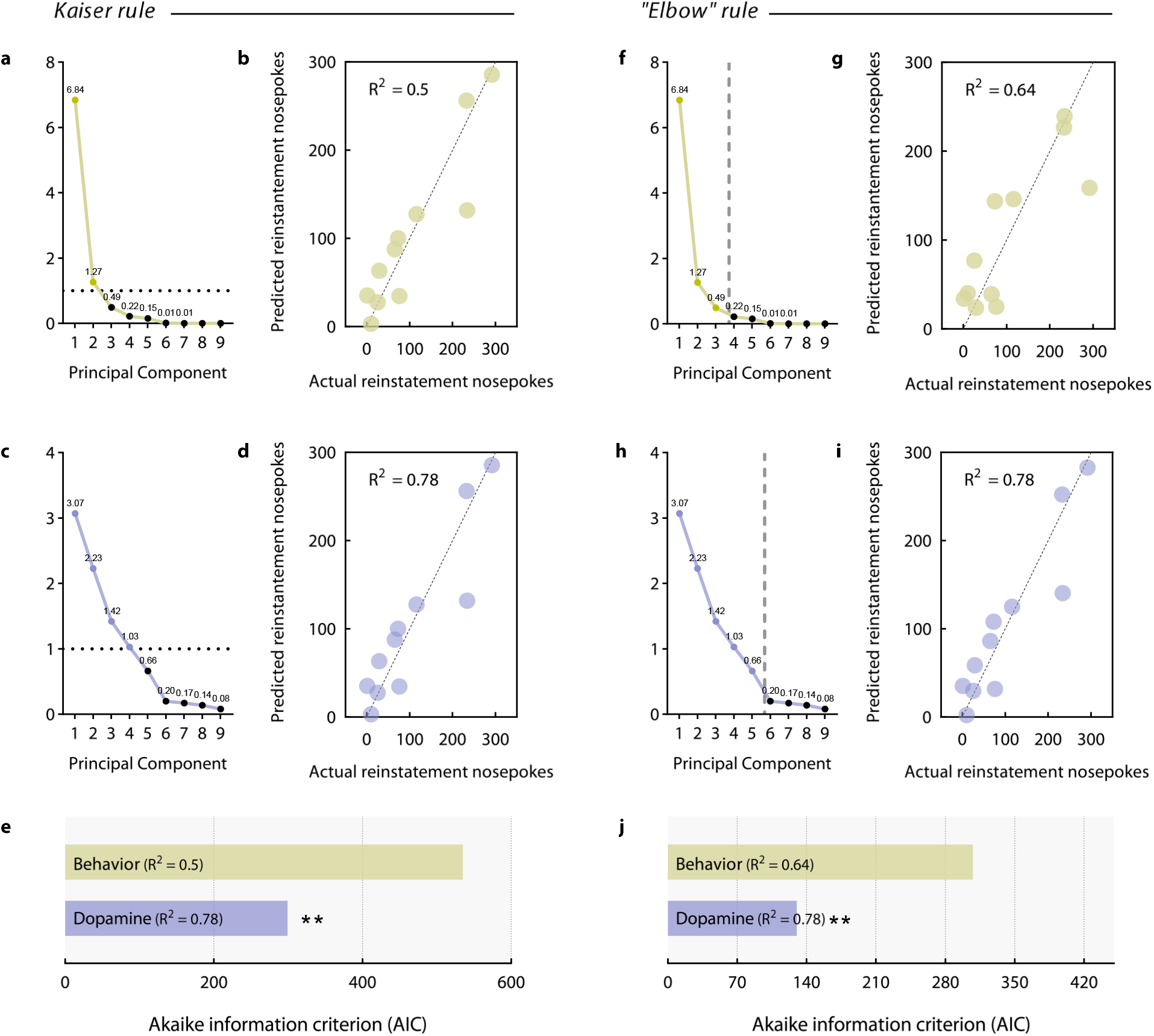
Different principal component selection methods for PCR lead to the same prediction about reinstatement behavior. (**a**) BE_PC_ eigenvalues of the principal components obtained after dimensionality reduction by PCA. The Kaiser rule selects all the principal components with an eigenvalue higher than 1. (**b**) Biplot of the observed and predicted reinstatement nosepoke values after running PCR with principal components BE_PC1_ and BE_PC2_ as covariates. (**c**) DA_PC_ eigenvalues of the principal components obtained after dimensionality reduction by PCA according to the Kaiser rule. (**d**) Biplot of the observed and predicted reinstatement nosepoke values after running PCR with principal components DA_PC1_, DA_PC2_, DA_PC3_ and DA_PC4_ as covariates. (**e**) AIC comparisons of the DA_PC_ and BE_PC_ MLR models of reinstatement following the Kaiser rule (ΔAIC = −235.9, \*\**p* = 0.002). (**f**) BE_PC_ eigenvalues of the principal components obtained after dimensionality reduction by PCA. The “elbow” rule selects all the principal components found before an apparent dip in the explanatory power of the next principal component. (**g**) Biplot of the observed and predicted reinstatement nosepoke values after running PCR with principal components BE_PC1_, BE_PC2_ and BE_PC3_ as covariates. (**h**) DAPC eigenvalues of the principal components obtained after dimensionality reduction by PCA according to the “elbow” rule. (**i**) Biplot of the observed and predicted reinstatement nosepoke values after running PCR with principal components DA_PC1_, DA_PC2_, DA_PC3_, DA_PC4_ and DA_PC5_ as covariates. (**j**) AIC comparisons of the DA_PC_ and BE_PC_ MLR models of reinstatement behavior following the “elbow” rule (ΔAIC = −130.2, \*\**p* = 0.004).

**Figure S7.**
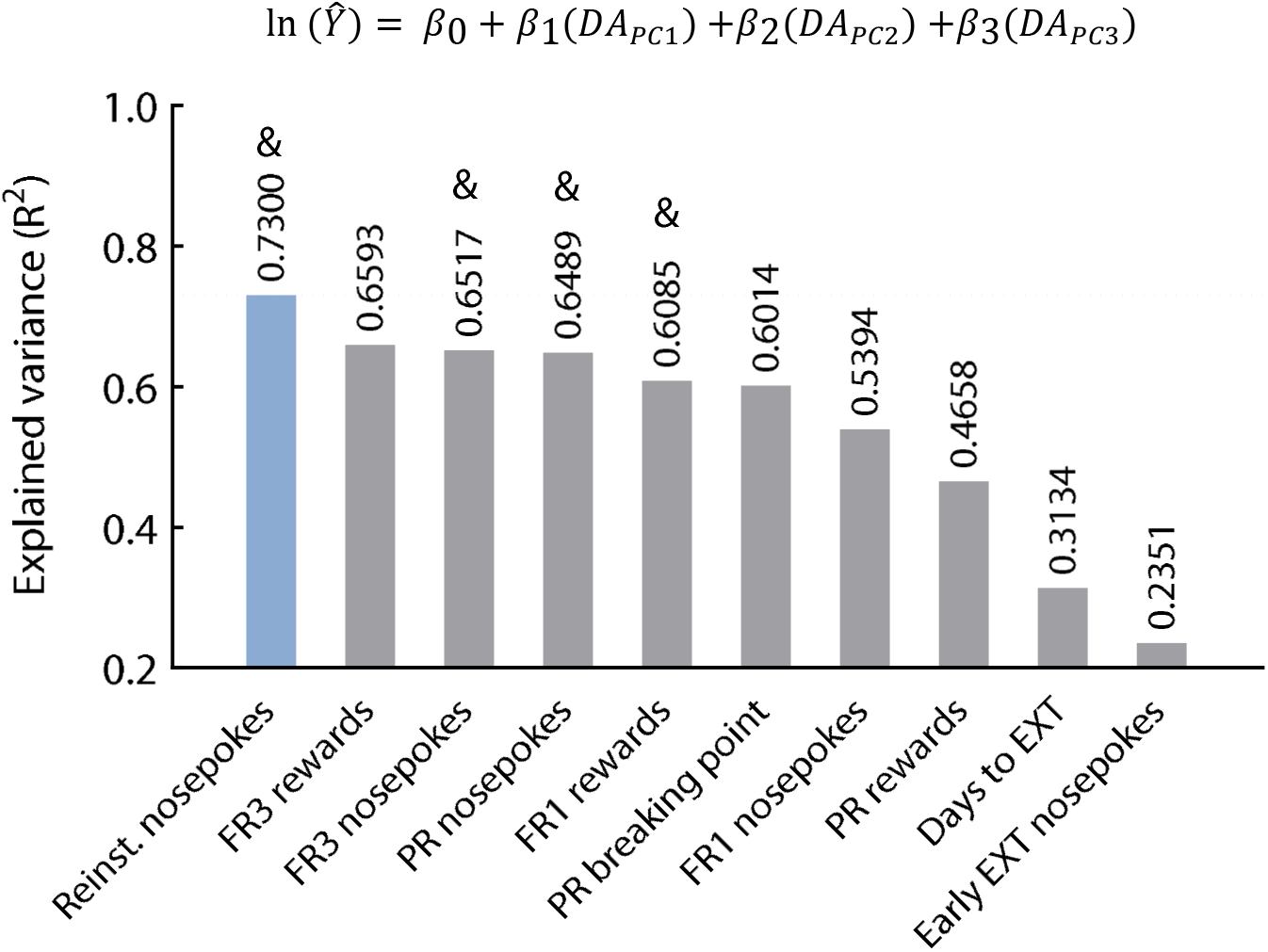
The DA_PC_ model predictions (R^2^) about other cocaine self-administration behavioral outcomes. Models in which all three DA_PC1_,DA_PC2_, and DA_PC3_ β-parameter estimates were significantly different from 0 are marked with &.

## Notes

### Competing Interest Statement

The authors have declared no competing interest.

